# An engineered ACE2 decoy broadly neutralizes Omicron subvariants and shows therapeutic effect in SARS-CoV-2-infected cynomolgus macaques

**DOI:** 10.1101/2022.12.29.522275

**Authors:** Emiko Urano, Yumi Itoh, Tatsuya Suzuki, Takanori Sasaki, Jun-ichi Kishikawa, Kanako Akamatsu, Yusuke Higuchi, Yusuke Sakai, Tomotaka Okamura, Shuya Mitoma, Fuminori Sugihara, Akira Takada, Mari Kimura, Mika Hirose, Tadahiro Sasaki, Ritsuko Koketsu, Shunya Tsuji, Shota Yanagida, Tatsuo Shioda, Eiji Hara, Satoaki Matoba, Yoshiharu Matsuura, Yasunari Kanda, Hisashi Arase, Masato Okada, Junichi Takagi, Takayuki Kato, Atsushi Hoshino, Yasuhiro Yasutomi, Akatsuki Saito, Toru Okamoto

## Abstract

The Omicron variant continuously evolves under the humoral immune pressure obtained by vaccination and SARS-CoV-2 infection and the resultant Omicron subvariants exhibit further immune evasion and antibody escape. Engineered ACE2 decoy composed of high-affinity ACE2 and IgG1 Fc domain is an alternative modality to neutralize SARS-CoV-2 and we previously reported its broad spectrum and therapeutic potential in rodent models. Here, we show that engineered ACE2 decoy retains the neutralization activity against Omicron subvariants including the currently emerging XBB and BQ.1 which completely evade antibodies in clinical use. The culture of SARS-CoV-2 under suboptimal concentration of neutralizing drugs generated SARS-CoV-2 mutants escaping wild-type ACE2 decoy and monoclonal antibodies, whereas no escape mutant emerged against engineered ACE2 decoy. As the efficient drug delivery to respiratory tract infection of SARS-CoV-2, inhalation of aerosolized decoy treated mice infected with SARS-CoV-2 at a 20-fold lower dose than the intravenous administration. Finally, engineered ACE2 decoy exhibited the therapeutic efficacy for COVID-19 in cynomolgus macaques. Collectively, these results indicate that engineered ACE2 decoy is the promising therapeutic strategy to overcome immune-evading SARS-CoV-2 variants and that liquid aerosol inhalation can be considered as a non-invasive approach to enhance efficacy in the treatment of COVID-19.

## INTRODUCTION

The Spike gene of the SARS-CoV-2 genome continuously mutates to increase the transmissibility and to escape neutralizing antibodies induced by vaccination and infection. Compared with the original Wuhan strain, Omicron (BA.1.1.529) possessed 26–32 mutations, three deletions, and one insertion in the spike protein, which achieved drastic immune evasion (1). Omicron subvariants emerges over time with additional mutations in the receptor binding domain (RBD) and exhibits further escapability. Through the global spread of BA.2 and BA.5, currently, as of November 2022, BA.5-derived BQ.1 and BA.2-derived XBB are predominant in several countries, including the United States, Singapore, and India (2-4).

We have previously developed engineered angiotensin-converting enzyme 2 (ACE2) containing mutations to enhance affinity toward SARS-CoV-2 spike, which achieved virus-neutralizing activity comparable to therapeutic monoclonal antibodies (5). The ACE2-based decoy is expected to be resistant to viral escape, because mutant virus escaping the ACE2 decoy would have limited capability to bind native ACE2 receptors on host cells and lose infectivity (6). In fact, engineered ACE2 decoy broadly neutralized not only SARS-CoV-2 variants including BA.1 and BA.2, but SARS-CoV-1 and other sarbecoviruses (1). In addition, engineered ACE2 decoy showed no signs of vulnerability to escape mutants when added at suboptimal concentrations during long-term SARS-CoV-2 culture (5), and deep mutational scanning (DMS) analysis showed that no single amino acid mutation in the spike RBD escaped engineered ACE2 decoy.

Inhaled administration is an effective and non-invasive drug delivery strategy and is highly preferred in lung diseases, such as asthma, chronic obstructive pulmonary disease, cystic fibrosis, and pneumonia (7). Compared with systemic administration, inhaled administration has several advantages, including rapid onset of action, sparing doses, higher concentrations delivered locally, and improved bioavailability (7, 8). As SARS-CoV-2 directly targets the respiratory tract, inhaled therapeutics could be a promising approach against COVID-19. Although no inhaled drugs are currently approved for COVID-19, some respiratory tract infections have been already treated through inhalation. For example, ribavirin is administered as an aerosolized mist via nasal and oral inhalation for the treatment of respiratory syncytial virus (9). Some neuraminidase inhibitors, such as zanamivir and laninamivir are used in the formulation of dry powder inhalation to treat influenza virus infections (10). To further maximize the therapeutic effect on COVID-19, inhaled administration of neutralizing drugs remains to be evaluated.

Here, we demonstrated that engineered ACE2 decoy efficiently neutralized BA.2, BA.5 and their sublineages including the extensively immune-evading BQ.1 and XBB. Intensive isolation of drug-resistant mutants using a replicating SARS-CoV-2 pseudovirus revealed that the high-affinity ACE2 decoy allowed no mutational escape even though escape mutants emerged in wild-type (WT) ACE2 decoy. As the therapeutic potential *in vivo*, the engineered ACE2 decoy treated Omicron subvariants, BA.2, BA.2.75 and BA.5 in rodent models. Moreover, inhaled aerosol administration was demonstrated to be effective at a twenty-fold lower dose, as compared with the standard intravenous injection. Finally, engineered ACE2 decoy conferred protection against infection with SARS-CoV-2 in a cynomolgus macaque model.

## RESULTS

### Safety of the engineered ACE2 decoy as a clinical drug candidate

We introduced several amino acid substitutions in human ACE2 protein to enhance the binding affinity toward the SARS-CoV-2 spike protein and to eliminate enzymatic activity for the risk of adverse effects due to over-conversion of angiotensin II to angiotensin 1-7 (Supplemental Figure 1A) (1). These mutations could activate the host immunity, which leads to the risk of anti-drug antibodies and immune cross-reactivity to the host tissues. To relieve this concern, the antigenicity of introduced mutations were evaluated with the T cell activation assay where overlapping peptides were incubated with peripheral blood mononuclear cells (PBMC) that contain both T cells and antigen presenting cells (APC) (11). Affinity-enhancing mutations from 3 ACE2 mutants (3N39v2, 3J113v2 and 3J320v2) and peptidase-dead mutations were analyzed with 10 different HLA donors-derived PBMCs (Supplemental Figure 1B-E). One peptide fragment containing L79F mutation in 3N39v2 stimulated T cells from multiple donors (Supplemental Figure 1C). We thus replaced L79F with the safe T92Q from 3J320v2, which removed N-glycan and preserved higher neutralization activity. The resultant ACE2 mutant has affinity-enhancing A25V/K31N/E35K/T92Q and peptidase-dead S128C/V343C, referred to as 3N39v4 (Supplemental Figure 1A).

Off-target binding is another major risk of unintended drug toxicity (12). Then, we assessed the binding proteins of 3N39v4-Fc using a membrane proteome array (13). Whereas the spike proteins of SARS-CoV-1 and SARS-CoV-2 bound to 3N39v4 as positive controls, no significant binding was detected in the human endogenous membrane proteins except Fcγreceptor (Supplemental Figure 1F). Next, we evaluated the cardiotoxicity of 3N39v4-Fc in human induced pluripotent stem cell-derived cardiomyocytes (hiPS-CMs), because development of many drug candidates has been delayed or cancelled due to their unexpected cardiotoxicity, such as Torsades de pointes (TdP) risk associated with QT interval prolongation (14). By recording of field potentials (FPs), treatment with 3N39v4-Fc did not cause prolongation of FP duration and early afterdepolarization (EAD), which is a trigger to initiate TdP (Supplemental Figure 2A-D). In addition, treatment with 3N39v4-Fc did not affect cardiac contractility, measured by imaging-based motion analysis. Taken together, we minimized predictable risk of drug toxicity in the final drug candidate, 3N39v4-Fc.

### Cryo-EM reveals three ACE2 decoys bind to trimeric spikes of SARS-CoV-2

Single-particle cryo-EM is a powerful technique for obtaining highly heterogeneous structures. We speculated that the WT ACE2 and 3N39v4 would bind to the RBDs of the spike proteins and affect the up/down states of these domains. First, 6 x His-tagged 3N39v4 and WT ACE2 were produced, and 6 x His-and foldon-tagged spike proteins were mixed to form complexes (Supplemental Figure 3A-C). As the results of image processing, several structures that differed in RBD states and ACE2 binding were obtained (Figure 1A, Supplemental Figure 4-6 and Supplemental Table 1). In WT-ACE2, 30% of the spike particles did not bind with ACE2 (1 up, no ACE2, Figure 1B), whereas only less than 10% of the particles did not bind with the 3N39v4. In addition, the distribution of the spike protein in which all three RBD occupied with three 3N39v4 was ∼28% which is much higher than that of WT ACE2 (∼18 %) (3 up, 3 ACE2, Figure 1B). It is expected that these distribution of the particle images in each class reflects the distribution of the corresponding state of the spike protein in the sample solution. Therefore, these results are consistent with the higher affinity of the 3N39v4 to the RBD of spike than WT ACE2 (5). The 3N39v4 binds strongly to the RBD when the domain adopts the up state. Once 3N39v4 is bound to the RBD, the 3N39v4 rarely dissociates from the domain and maintains the up state of the domain. Thus, the distribution of the particles shifted toward a higher number of ACE2 binding sites than that of the WT ACE2.

**Figure 1.**
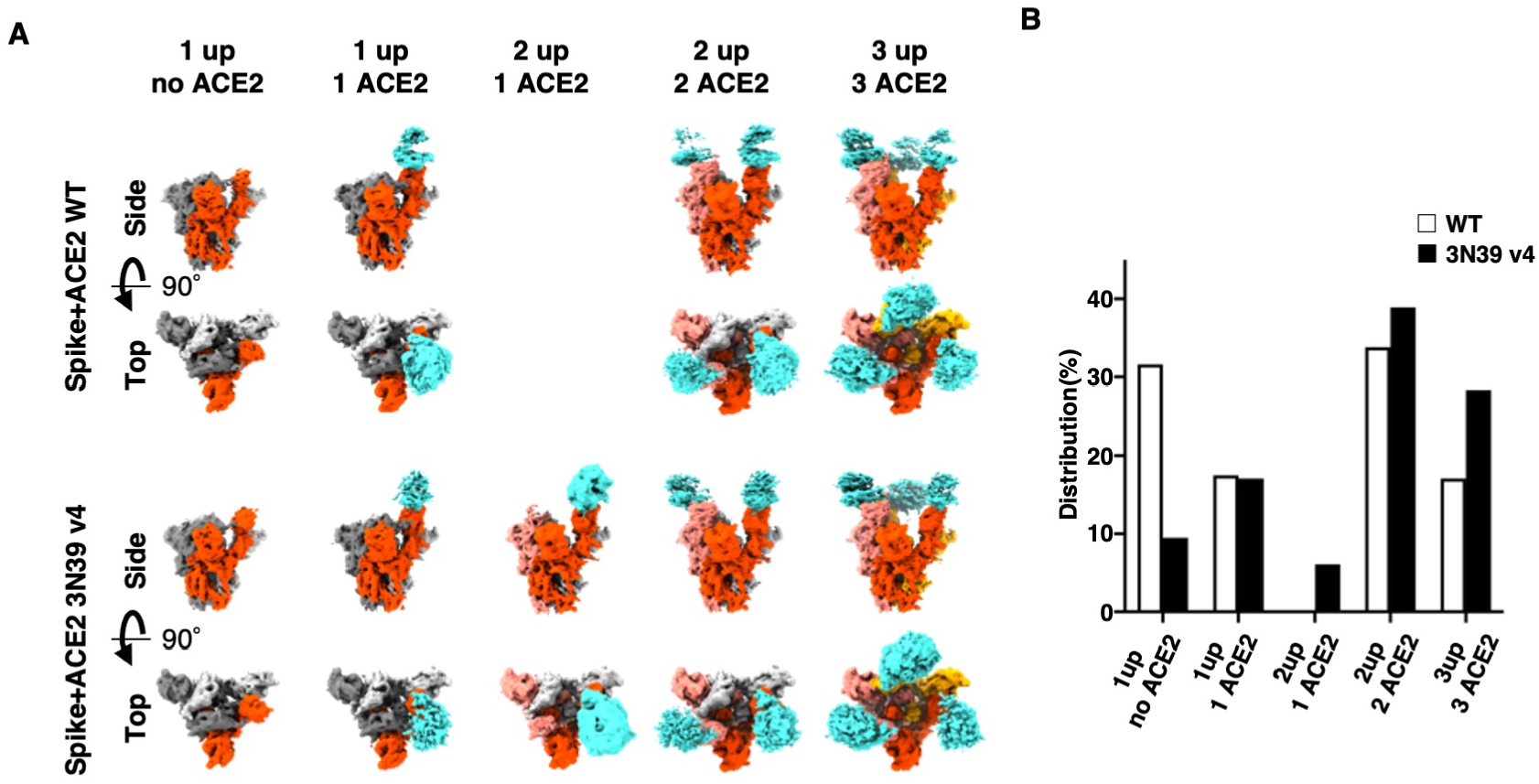
Structural insights of interaction between ACE2 decoy and SARS-CoV-2 spike. (A) Obtained maps from the image processing of WT ACE2-spike and 3N39v4-spike complexes. The maps were gaussian filtered using UCSF chimera tools. The protomer in up or down states are colored red or gray colors, respectively; ACE2 and 3N39v4 are colored by cyan. (B) Dstribution of particles used final 3D classification. The workflow of image processing is summarized in Supplemental Figure 4 and 5.

### Engineered ACE2 decoy counteracts the emergence of escape mutations

The main advantage of engineered ACE2 decoy is that it prevents the emergence of escape mutants. SARS-CoV-2 mutations that confer escape from therapeutic antibodies can arise especially in immunocompromised patients with prolonged infection and repeated drug administration. Such viral evasion may contribute to the emergence of global variants of concern (15,16). We previously reported that continuous viral passages (up to 15 times) of SARS-CoV-2 under serial concentrations of engineered ACE2 decoy produced no escape mutants in contrast to the emergence of antibody escape at 4th passage (5). In this study, we used a replication-competent VSV possessing the SARS-CoV-2 Spike gene (VSV-SARS-CoV-2-S) which exhibited higher mutation rate due to the lack of the proof-reading coronavirus exonuclease (17) (Figure 2A). VSV-SARS-CoV-2-S was incubated with a series of WT ACE2-Fc, 3N39v4-Fc, or bebtelovimab (18) (Figure 2B). After seven passages, VSV-SARS-CoV-2-S cultured with WT ACE2-Fc or bebtelovimab conferred drug resistance (Figure 2C). WT ACE2-resistant VSV-SARS-CoV-2-S also exhibited some tolerance to 3N39v4-Fc, but still remained functionally sensitive due to higher binding affinity in baseline (Figure 2D). This mutant obtained N370D and Y489H in the RBD and T791A in the S2 domain (Figure 2E). In the neutralization assay for pseudoviruses possessing N370D, Y489H, and/or T791A, the N370D mutant was neutralized similarly to parental virus, but both Y489H and N370D/Y489H/T791A had lower sensitivity to WT ACE2-Fc and 3N39v4-Fc (Figure 2F), suggesting that Y489H mutation is responsible for reduced neutralization by ACE2 decoys. To understand the mechanism of Y489H-mediated escape from ACE2 decoys, we characterize the Y489H mutation on the binding affinity, the spike protein expression, and infectivity. The binding of WT ACE2-Fc and spike-expressing cells was evaluated by flow cytometry. Consistent with the neutralization assay, Y489H and N370D/Y489H/T791A had substantially lower binding to WT ACE2-Fc, compared with N370D and parental spike (Figure 2G). In contrast, spike protein expression on the cell surface was upregulated in Y489H (Figure 2H), which resulted in relatively preserved infectivity (Figure 2I). The level of spike protein expression is one of the determinants of viral infectivity. The Y489H mutation compromised the binding to ACE2 but compensatively increased spike protein even with the structurally fragile N370D, which collectively lead to functional infectivity with the escape from WT ACE2-Fc decoy.

**Figure 2.**
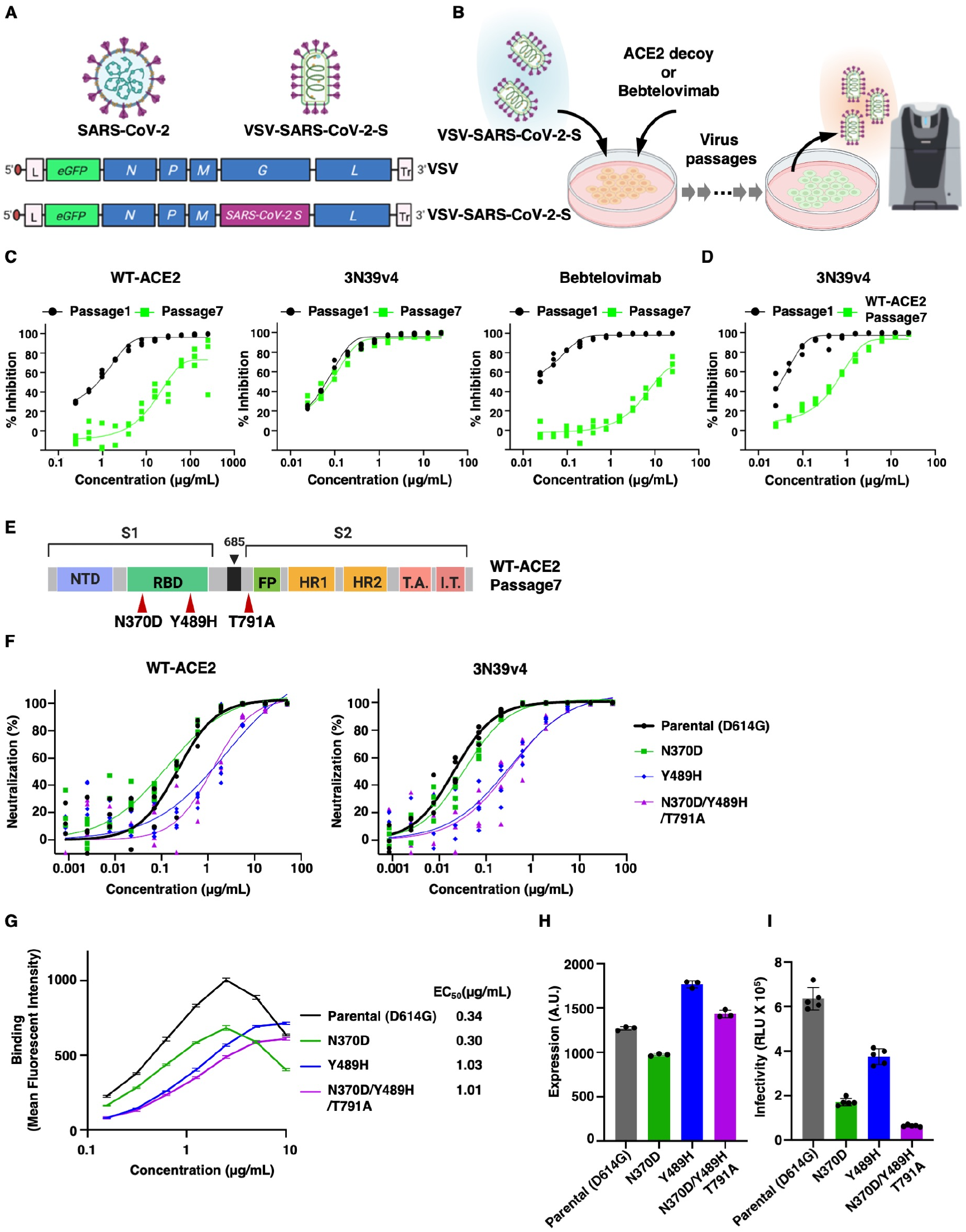
Engineered ACE2 decoy allowed no emergence of escape mutant viruses. **(A)** Schematic construct of VSV possessing the SARS-CoV-2 spike protein. **(B)** VSV-SARS-CoV-2-S was cultured with ACE2 decoy or bebtelovimab. After imaging cytometry analysis, supernatants with 50% reduction in neutralizing activity were repeatedly passaged in naïve Huh7/ACE2 cells. **(C)** Neutralization efficacy is shown for WT ACE2-Fc (left), 3N39v4-Fc (middle), and bebtelovimab (right) using VSV-SARS-CoV-2-S at 1st and 7th passages (n = 3 technical replicates). **(D)** Neutralization efficiency using VSV-SARS-CoV-2-S passaged 7th under WT ACE2-Fc is shown using 3N39v4-Fc (n = 3 technical replicates). **(E)** Scheme of SARS-CoV-2 spike protein is illustrated with three mutations emerging in the WT ACE2-Fc resistant virus. **(F)** Neutralization efficacy was measured in 293T/ACE2 cells for WT ACE2-Fc or 3N39v4-Fc against the indicated escape candidates (n = 4 technical replicates). **(G)** Avid binding was measured by flow cytometry of WT ACE2-Fc to cells expressing HA-tagged full-length spike of indicated escape candidates. EC_50_ was indicated for each mutation (n = 3 technical replicates). **(H)** Spike protein expression was measured by flow cytometry using anti-HA antibody (n = 3 technical replicates). **(I)** The infectivity of pseudotyped lentivirus carrying spike protein was evaluated with firefly luciferase-reporter system (n = 5 technical replicates). Data are presented as mean ± s.e.m..

### Engineered ACE2 decoy confers protection against infection with SARS-CoV-2 BA.4, **BA.5**, and BA.2.75

We examined the efficacy of 3N39v4-Fc in Omicron subvariants *in vitro* and *in vivo*. In the pseudovirus experiment, 3N39v4-Fc neutralized BA.2, BA.4/5 and these derivatives including BA.2.12.1, BA.2.75, and currently spreading XBB and BQ.1 (Figure 3A). In contrast, both XBB and BQ.1 evaded bebtelovimab that was the only monoclonal antibody effective to both BA.2 and BA.4/5 in clinical use (19) (Figure 3B). Consistently, authentic viruses, including BA.1, BA1.18, BA.2, BA.2.75, XE, BA.4, and BA.5, were similarly neutralized by 3N39v4-Fc (Figure 3C).

**Figure 3.**
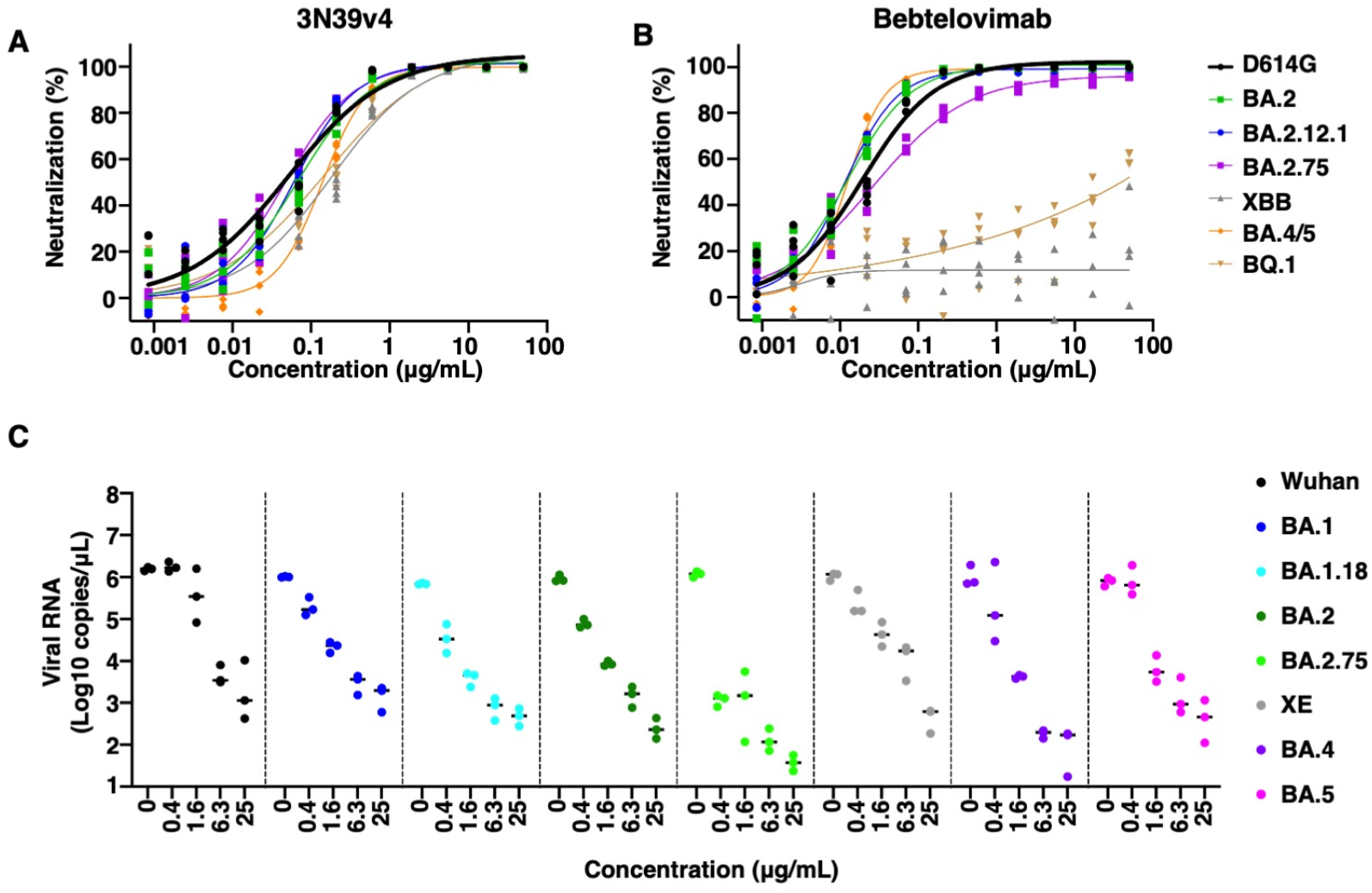
Engineered ACE2 decoy neutralizes a broad range of Omicron subvariants. **(A, B)** Neutralization efficacy in 293T/ACE2 cells is shown for 3N39v4-Fc (A) or bebtelovimab (B) against Omicron subvariants pseudoviruses (n = 4 technical replicates). **(C)** Sensitivity of 3N39v4 to the Wuhan and Omicron-derived variants was compared using each infectious virus in VeroE6/TMPRSS2 cells. RNA copy number was analyzed using qRT-PCR against nucleocapsid. Data are presented as mean of n = 3 technical replicates.

We also tested efficacy of 3N39v4-Fc to Omicron subvariants in a hamster model (Figure 4A). Hamsters were infected with 1 × 10^6^ median tissue culture infectious dose (TCID_50_) of BA.4, or BA.5 variants and followed by the intraperitoneal injection of 3N39v4-Fc at 2-h intervals. After 5 days, viral RNA genome in the lungs was significantly reduced by treatment with 3N39v4-Fc (Figure 4B and 4C). The expression of genes encoding inflammatory cytokines, such as Il6, Cxcl10, Ccl5, and Ccl3 was attenuated by 3N39v4-Fc treatment in hamsters infected with BA.4 (Figure 4D) or BA.5 (Figure 4E) variants. These data suggest that 3N39v4-Fc could be used as a therapeutic agent for continuously emerging Omicron subvariants.

**Figure 4.**
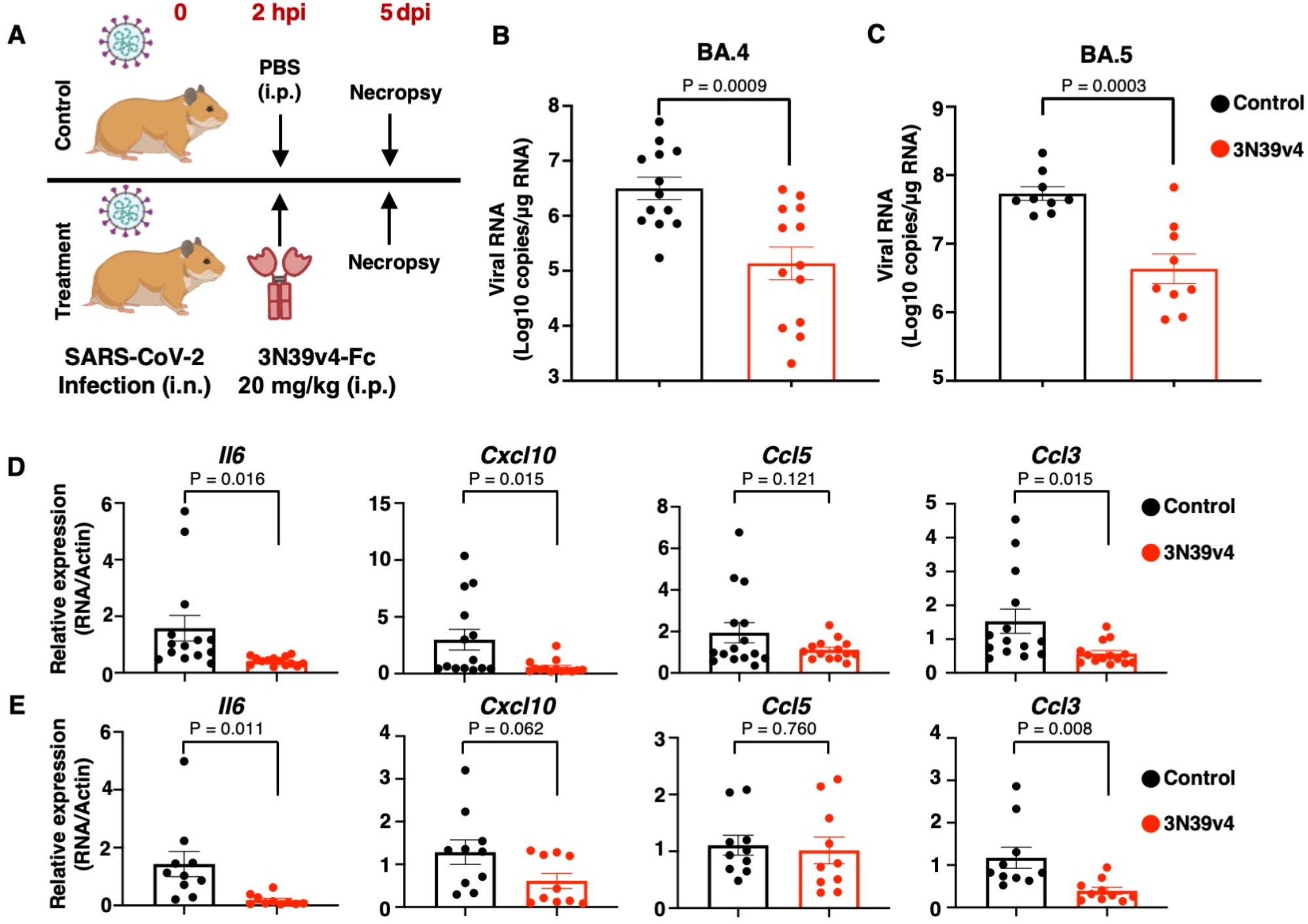
Engineered ACE2 decoy confers therapeutic efficacy against Omicron subvariants *in vivo*. **A)** Experimental design for establishment of a hamster model of SARS-CoV-2. Syrian hamsters were intranasally inoculated with SARS-CoV-2 (BA.4 or BA.5) using 1.0 × 10^5^ TCID_50_ (in 60 μL). **(B, C)** Viral RNA load in the lungs of control (BA4; *n* = 12, BA.5; *n* = 8) or 3N39v4-Fc-treated (BA4; *n* = 12, BA.5; *n* = 8) hamsters at 5 dpi. **(D, E)** mRNA expression of inflammatory cytokines and chemokines in the lungs from hamsters infected with BA.4 (D) or BA.5 (E) at 5 dpi. Data are represented as mean ± s.e.m. *P* values were determined using two-tailed, unpaired parametric *t*-tests. hpi, hours post infection; dpi, days post infection; i.n., intranasally; i.p., intraperitoneally.

### Biodistribution of engineered ACE2 decoy administrated via inhaled routes

Systemic delivery of neutralizing drugs is inefficient to achieve optimal drug concentration for respiratory tract infection. In contrast, liquid aerosol inhalation is the direct delivery to the infection site and expected to be more effective. To evaluate the efficacy of 3N39v4-Fc inhalation, we first analyzed the pharmacokinetics of inhaled 3N39v4-Fc in mice with ^89^Zirconium (Zr)-labelling and positron emission tomography (PET) imaging (20). PET imaging after intravenous administration of ^89^Zr-labelled 3N39v4-Fc at a dose of 400 μg/body showed radioactivity of the blood pool rapidly decreased and constant uptake in the liver (Figure 5A-C). This biodistribution is the similar pattern to the monoclonal antibody distribution (21) and there was no other abnormal uptake that indicated the possibility of off-target binding and the risk of unintended adverse effects. In contrast, inhaled administration of ^89^Zr-labelled 3N39v4-Fc at a dose of 80 μg/body revealed that dominant uptake in lungs diminished over time and obvious distribution was observed in the stomach 24-h after inhalation, indicating that inhaled 3N39v4-Fc was mainly drained into the digestive tract by mucociliary clearance (22) (Figure 5D-F).

**Figure 5.**
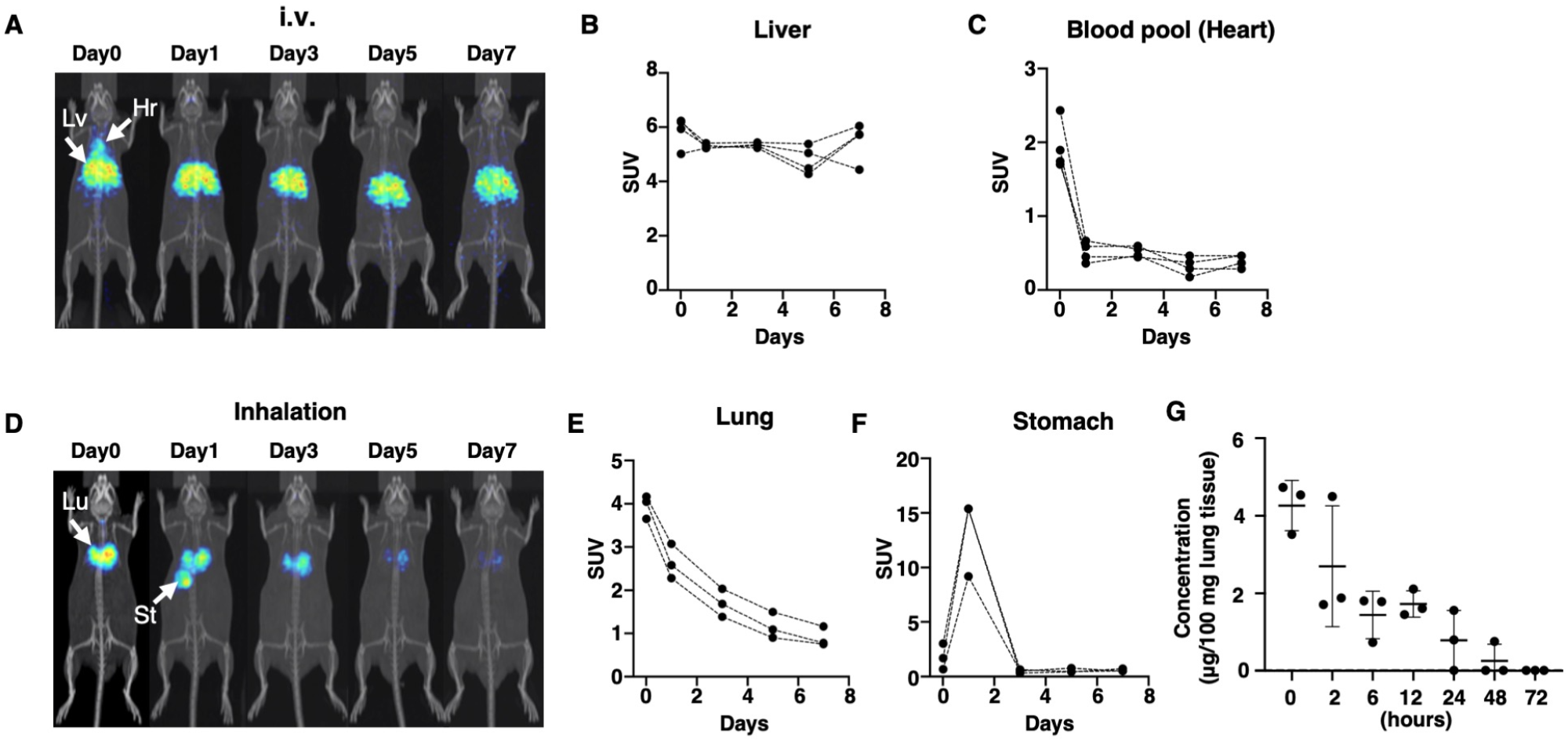
Enhanced lung distribution of engineered ACE2 decoy in the inhaled administration. **(A, D)** Distribution of ^89^Zr -labeled 3N39v4-Fc after intravenous administration (A) or after inhalation (D) in mice. Coronal maximum intensity projection (MIP) PET images of ^89^Zr -labeled 3N39v4 was visualized at indicated time points. Hr, heart; Lv, liver. Lu, lung; ST, stomach. **(B, C)** Time-activity curves of 3N39v4-Fc in individual organs in the liver (B) and heart (C) were quantified (n=4). **(E, F)** Time-activity curves of 3N39v4-Fc in the lung (E) and stomach (F) were quantified (n=3). **(G)** The concentration of 3N39v4-Fc in the lung were determined by ELISA at indicated time points. Data are presented as mean ± s.d..

We further evaluated the condition of inhaled 3N39v4-Fc in the lung. The lung lysate was separated with SDS-PAGE and the radioactivity was measured for pre-loading lysate and 3N39v4-Fc segment after electrophoresis (Supplemental Figure 7A). The fraction of Intact ^89^Zr-labelled 3N39v4-Fc in the total ^89^Zr signal was 30.1%, 24.6, and 14.2% in the non-reduced SDS-PAGE on days 1, 4, and 7, respectively (Supplemental Figure 7B). Based on these results, approximately 40% of inhaled 3N39v4-Fc was washed out mainly into the digestive tract, 30% was degraded or aggregated in the lung and another 30% remained non-degraded dimer form in day 1. Similar result for non-degraded 3N39v4-Fc was observed by ELISA which captured 3N39v4-Fc by anti-Fc antibody and detected it by anti-ACE2 antibody to analyze non-degraded 3N39v4-Fc (Figure 5G)

### Inhaled administration efficiently treats SARS-CoV-2 infection

To compare the anti-viral effects of 3N39v4-Fc through intravenous or inhaled administration, Balb/c mice were intranasally infected with 1 × 10^4^ TCID_50_ of mouse-adapted SARS-CoV-2 (SARS-CoV-2 MA) (23), and 3N39v4-Fc was administered via intravenous or inhaled route 24 h after inoculation (Figure 6A). Twenty of the 22 mice treated with PBS inhalation as a control died within 7 days after inoculation with more than 20% body weight loss (Figure 6B). In the inhaled treatment group, mice survival rate was improved in a dose-dependent manner and 14 of 25 mice survived SARS-CoV-2 MA infection with ∼15% body weight loss in 1 mg/kg of 3N39v4-Fc inhalation, which was equivalent efficacy to 20 mg/kg of 3N39v4-Fc intravenous administration (Figure 6C). Viral RNA genome, virus titer and expression of inflammatory cytokine, IL-6, in the lungs were suppressed in mice administered by 0.25 mg/kg and 1.0 mg/kg via inhaled routes and 20 mg/kg and 5 mg/kg, but not 1 mg/kg, via intravenous routes (Figure 6D-F). We further confirmed that 1 mg/kg of 3N39v4-Fc inhalation had the therapeutic efficacy equivalent to 20 mg/kg of intravenous administration in hamsters infected with BA.5 and BA.2.75 variants (Figure 6G-J,). Collectively, these data suggest that inhalation of 3N39v4-Fc can effectively treat SARS-CoV-2 infection with dose-sparing benefits up to 20-fold in rodent models.

**Figure 6.**
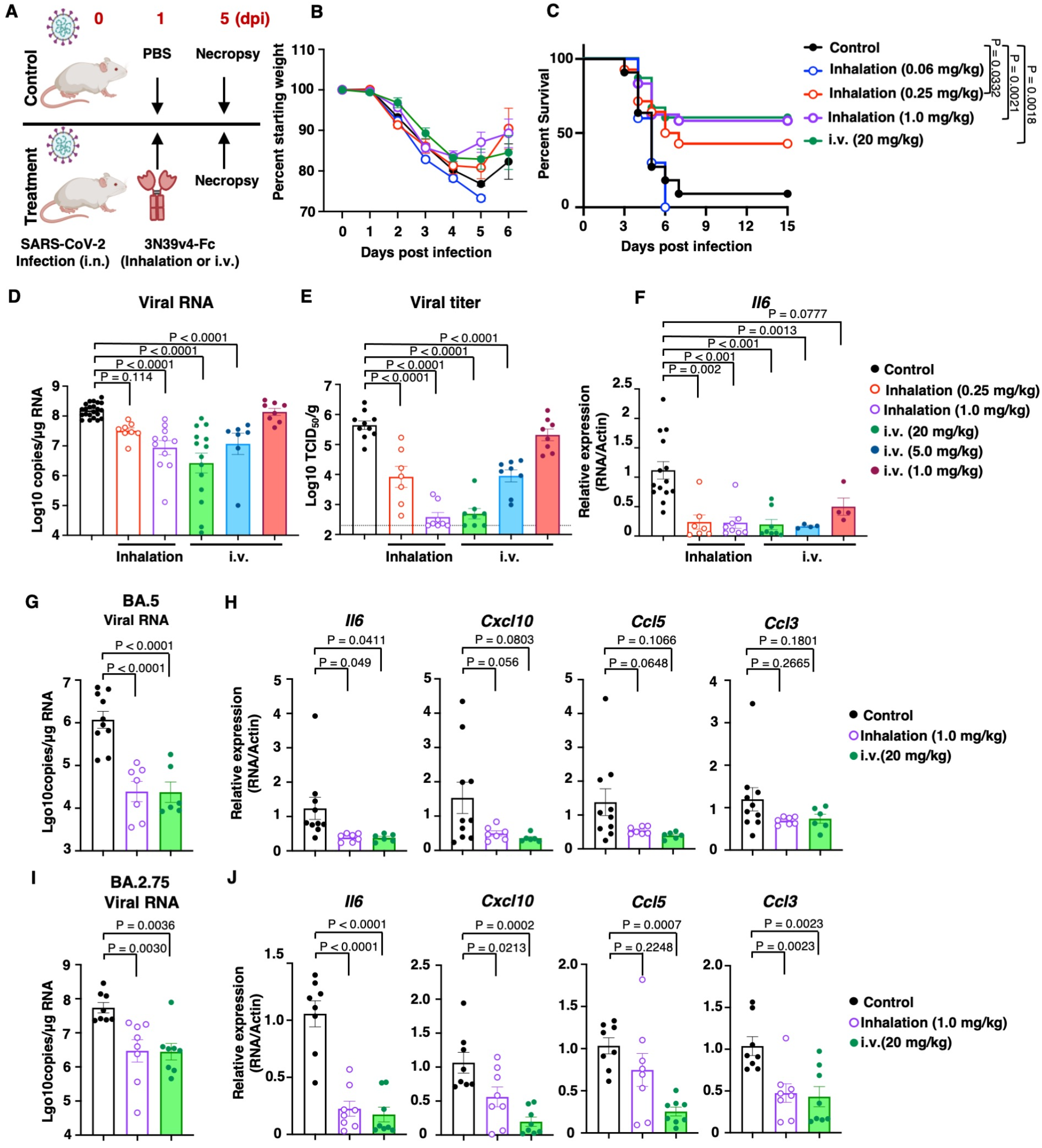
Inhaled administration enhances efficacy of engineered ACE2 decoy. **(A)** Experimental design for establishment of a mouse model using mouse-adapted SARS-CoV-2 (MA10). Balb/c mice infected intranasally (i.n.) with MA10 at 1.0 × 10^4^ TCID_50_ (in 20 μL). **(B, C)** The body weight and a survival curve (C) are shown for Balb/c mice (control, *n* = 22; inhalation: 0.06 mg/kg, *n* = 10; inhalation: 0.25 mg/kg, *n* = 14; inhalation: 1.0 mg/kg, *n* = 25, and i.v: 20 mg/kg, *n* = 16). **(D-F)** Viral RNA (D), infectious viral titer (E) and mRNA expression of Il6 (F) in the lungs of control or 3N39v4-Fc-treated mice at 5 dpi. **(G, H)** Syrian hamsters were intranasally inoculated with SARS-CoV-2 (BA.5) at 1.0 × 10^6^ TCID_50_ (in 60 μL) and treated with PBS via inhaled routes (Control) or 3N39v4-Fc via inhaled routes (inhalation) or intravenous routes (i.v.) (*n* = 8) at 24 hours post infection (hpi). Viral RNA load (G), and mRNA expression of Il6, Cxcl10, Ccl5 and Ccl3 (H) in the lungs of control or 3N39v4-treated hamsters at 5 dpi. **(I, J)** Syrian hamsters were intranasally inoculated with SARS-CoV-2 (BA.2.75) at 1.0 × 10^6^ TCID_50_ (in 60 μL) and treated with PBS via inhaled routes (Control) or 3N39v4-Fc via inhaled routes (inhalation) or intravenous routes (i.v.) (*n* = 8) at 24 hpi. Viral RNA load (I), and mRNA expression of Il6, Cxcl10, Ccl5 and Ccl3 (J) in the lungs of control or 3N39v4-treated hamsters at 5 dpi. Data are presented as mean ± s.e.m. *P* values were determined using one-way ANOVA, followed by Dunnett’s multiple comparison test (D-J). dpi, days post infection; i.n., intranasally; i.v., intravenously; i.p., intraperitoneally.

### Engineered ACE2 decoy ameliorates pneumonia induced by SARS-CoV-2 infection in a cynomolgus macaque model

To test the therapeutic effect of 3N39v4-Fc in a non-human primate model, we employed a cynomolgus macaque (CM) and the Delta variant that induces apparent lung pathology as compared with other variants (24). Nine CMs were inoculated with 1 × 10^6^ PFU of Delta variant via a combination of intratracheal (i.t.) and intranasal (i.n.) routes at 0 dpi. Four animals (Control group) were intravenously injected with PBS as a control and five animals (Treatment group) were treated with 50 mg/kg of 3N39v4-Fc via intravenous administration at 1 dpi (Figure 7A). The half-life of 3N39v4-Fc was approximately 60 h and the blood concentration was preserved more than 20 µg/mL at 7 dpi (Figure 7B). The Control group showed multilobular ground glass opacity on the computed tomography (CT) imaging, in contrast, lung abnormalities were limited in the Treatment group (Figure 7C). Notably, at 3 and 5 dpi, the amount of infectious virus in the nasal swab samples of the Treatment group was significantly reduced to undetectable levels (Figure 7D). While the difference between Control group and Treatment group was not significant, the Treatment group had undetectable level of infectious virus in the pharynx swab at 3, 5 and 7 dpi (Supplemental Figure 7A). There were no significant differences in the body temperature, blood parameters, body weight, or clinical score (Supplemental Figure 8B-H).

**Figure 7.**
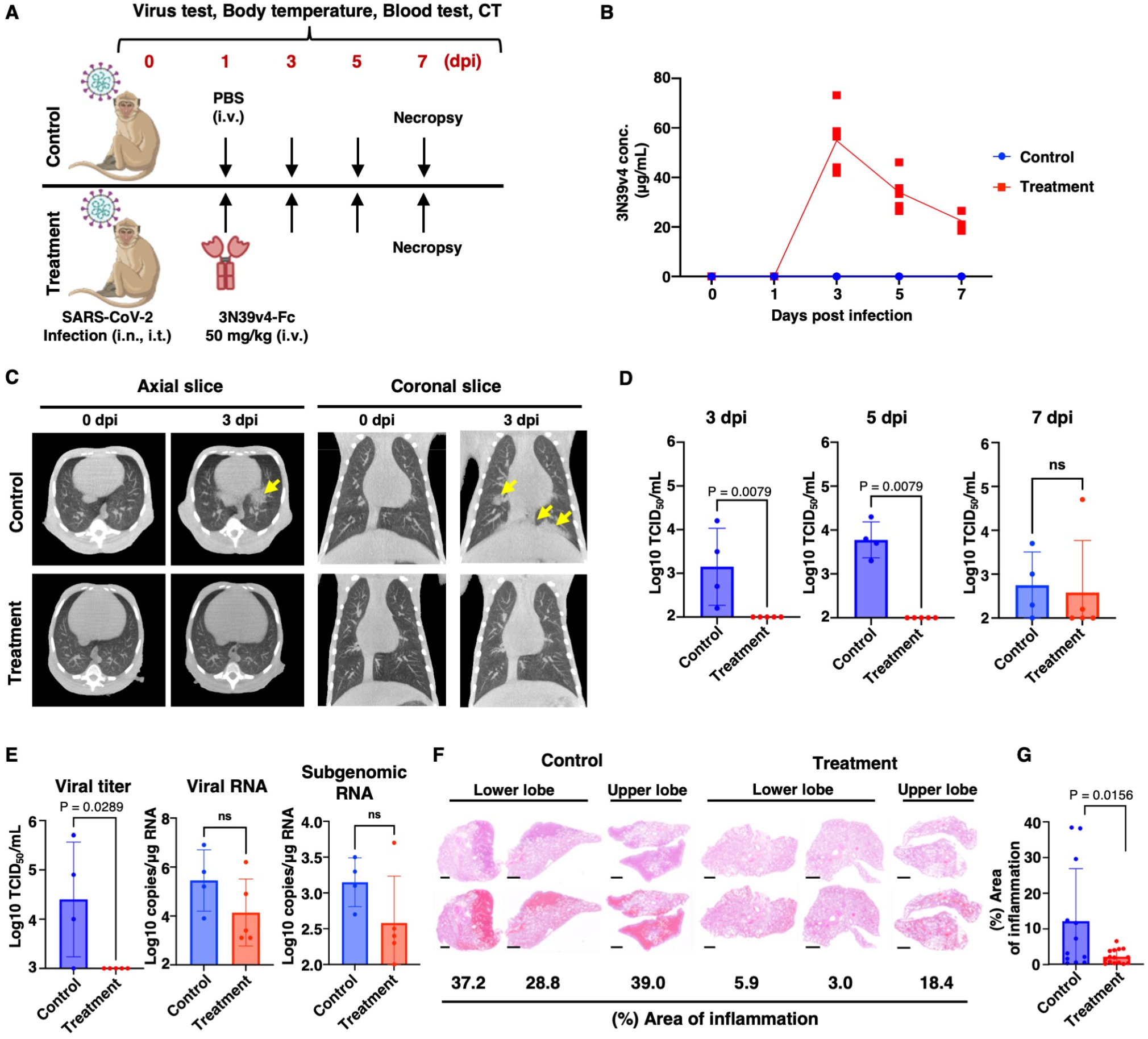
High affinity ACE2 decoy exhibits therapeutic effect in a cynomolgus macaque model. **(A)** Schematic procedure of in vivo experiments in a monkey model. Nine male cynomolgus monkeys were inoculated with SARS-CoV-2 Delta isolates via a combination of intratracheal and intranasal routes, with 1 × 10^6^ TCID_50_ virus. At 1 dpi. Four monkeys (Control group) were intravenously inoculated with PBS, and five monkeys (Treatment group) received intravenous 3N39v4-Fc inoculation. **(B)** 3N39v4-Fc concentration in monkey plasma samples. concentration of 3N39v4-Fc in the monkey plasma samples was measured at the indicated time points. **(C)** Pulmonary inflammation in SARS-CoV-2–infected CMs. Chest CT images were obtained during the experimental period. The red marker indicates a representative inflammation image of pneumonia. **(D)** Viral shedding in nasal swab samples. After SARS-CoV-2 inoculation, nasal swab samples were collected at the indicated time points. Infectious viral titers in the swab samples were determined based on TCID_50_ using VeroE6/TMPRSS2 cells. Differences between Control and Treatment groups were compared. **(E)** Viral replication in the lung. At the time of euthanasia, lung tissues were minced to measure the levels of viral titer (left), viral RNA (center), and subgenomic RNA (right). **(F)** The lung tissue sections of the animals in the Control and Treatment groups were stained with H&E, and the ratio of the inflammatory area to the total lung area was calculated. HE-stained sections (upper column) and HE-stained sections with red highlight on inflammatory area (lower column) are shown. Scale bar: 2.5 mm. **(G)** Data are presented as average ratio and s. d. of each group, and each dot represents the ratio of each tissue section.

Monkeys were euthanized at 7 dpi to examine the viral titer, viral RNA, and subgenomic RNA levels in the lungs. Although we found modest difference in the levels of viral RNA and subgenomic RNA between the two groups, the viral titer in all animals in the Treatment group was significantly suppressed to the limit of detection, as compared with animals in the Control group (Figure 7E). These observations demonstrate that the 3N39v4 significantly suppresses infectious virus expansion in the lungs. Furthermore, histopathological examination showed severe pneumonia characterized by widespread infiltration of inflammatory cells and alveolar hemorrhage in the Control group, whereas these pathological alterations were significantly mitigated in the Treatment group (Figure 7F, G).

## DISCUSSION

Engineered ACE2 decoy has neutralizing activity comparable to monoclonal antibodies and broad spectrum targeting all major SARS-CoV-2 variants and even some sarbecoviruses. Decoy approaches that closely mimic viral receptors could overcome mutational escape, because viral resistance through mutation cannot develop without the concomitant loss of infectivity. As hypothesized, even highly mutating VSV-SARS-CoV-2-S failed to escape engineered ACE2 decoy in the treatment with suboptimal drug concentrations. However, some mutants emerged in WT ACE2-Fc treatment. Historically, this strategy has been applied to HIV-1 treatment that requires long-term viral control. Initial attempts to design decoys simply used a soluble CD4, the main receptor for HIV-1. However, WT soluble CD4 proved ineffective in human patients. Similar to the case of WT ACE2-Fc, isolated HIV-1 mutants were relatively insensitive to soluble CD4 neutralization without apparent loss of viral fitness (25,26). The strength of infection and neutralization can be determined by binding affinity and avidity effect. In the case of soluble CD4 monomer and ACE2-Fc dimer, affinity-lowering viral mutations reduce the sensitivity to decoys, but infectivity can be compensated by the avidity effect due to increased structural stability and multiple interaction (27). One approach to counteract this kind of mutation is to increase avidity with the multivalency of decoys (6,27). In the case of HIV-1, nanoparticle carrying CD4 decoy clusters effectively controlled HIV-1 replication without the emergence of escape mutation (26). Another approach is the engineering of high affinity decoys. We demonstrated that affinity-enhancing approach can achieve the escape-free virus neutralization in SARS-CoV-2 infection, and this approach is expected to be appliable for other viral infection diseases.

Inhaled administration is expected to be an effective delivery of therapeutic agents for respiratory diseases. In the present study, we showed that inhalation of 1.0 mg/kg 3N39v4-Fc had similar therapeutic effects to the intravenous dose of 20 mg/kg on the survival rate, suppression of viral load, and expression of inflammatory cytokines, indicating a 20-fold dose sparing benefit. Previous reports also demonstrated that the antiviral effect of 0.35 mg/kg remdesivir inhalation was similar to 10 mg/kg intravenous remdesivir (28), and that inhaled 100–200 μg of salbutamol was therapeutically equivalent to 2-4 mg of oral salbutamol (29). In case of COVID-19, several studies also demonstrated the therapeutic potential of inhaled monoclonal antibodies (30) and the ACE2 decoy (31) in animal models. In addition, we report, for the first time, the detailed pharmacokinetics of inhaled biologics with ^89^Zr-labelling and PET imaging. One of the challenges of inhalation is the fast clearance, less than 24 h half-life in the case of monoclonal antibodies in human (32). Inhaled drugs can be cleared from pulmonary tissues through various elimination pathways, such as coughing, mucociliary transport, phagocytosis by alveolar macrophages, and translocation into the cells and blood (33). The PET imaging in this study indicated that 40% of inhaled engineered ACE2 was transported into the digestive tract, 30% was degraded in the lung tissue and another 30% was expected to be active for virus neutralization 24 h after inhalation. Against this rapid clearance, several approaches have been reported to improve their structural integrity and bioavailability in pulmonary tissues, including Fc engineering, PEGylation, addition of excipients, and muco-trapping (33-35). Fusion of the IgG1 Fc domain prevents the degradation of proteins through binding to the neonatal Fc receptor (FcRn), expressed in the upper and central airways of the lung. In the pinocytosis-mediated IgG internalization and endosomal sorting, Fc can be captured by FcRn at endosomal pH (5.5–6.0) and directed to the plasma membrane. This sorting prevents the degradation in the endolysosome and pH-dependent binding affinity leads to the release from FcRn at extracellular pH 7.4 and recycled back into the airway (36). Although it is important to consider the optimal formulation to keep bioavailability during the manufacture or production of aerosols for inhalation, bioactive proteins possessing Fc fragments have been demonstrated to be delivered to the lungs via inhalation using a nebulizer in monkeys (37). In addition, the dry powder of monoclonal antibodies was delivered to the lungs with high bioavailability (38). These findings suggest that inhalation of 3N39v4-Fc has potential for practical use in COVID-19 treatment. The IgA-Fc-fused nanobody exhibited more potent inhibitory activity than the monomeric nanobody and IgG-Fc fusion against SARS-CoV-2 because of improvement of binding activity with the RBD of SARS-CoV-2, and inhaled IgM antibodies have long-term retention in the nasal cavity and lungs (33,39-41). These strategies could improve the efficacy of ACE2 decoy development in the future. Moreover, although a single dose of the 3N39v4 showed therapeutic effects in our rodent models, increasing the frequency of administration may demonstrate higher efficacy.

Based on the successful treatment of COVID-19 in rodent models, we tested the therapeutic effect of 3N39v4-Fc in a cynomolgus macaque model. Intravenous administration of 3N39v4-Fc significantly decreased not only the lung pathology but the number of infectious viruses in the nasal swabs, suggesting that the use of 3N39v4-Fc can decrease the risk of transmission from infected individuals. Although we used a fixed dose of the 3N39v4-Fc (50 mg/kg) in this study, a lower dose should be tested in a future study. Our preliminary data using one monkey demonstrated that a lower dose (20 mg/kg) had a therapeutic effect against SARS-CoV-2 infection. In addition, the efficacy of inhalation should be examined. To reduce the risk of adverse effects, we previously eliminated the ACE2 peptidase activity by introducing the disulfide-bond to close the angiotensin II binding pocket. The peptidase dead modification prevents the over-conversion of angiotensin II to angiotensin 1-7 and subsequent possible hemodynamic dysregulation. The other main issues related to drug toxicity are immunogenicity and off-target binding to endogenous proteins. With T cell assays from 10 different HLA donors, L79F substitution induced obvious T cell response, thus we replaced it with the safe T92Q substitution which enhances the binding affinity by removing N-glycan. Screening for off-target with membrane proteome array indicated no apparent endogenous binding target except Fc receptor. After these risk-predicting assays, we decided 3N39v4-Fc as the final candidate decoy.

In summary, engineered ACE2 decoy, 3N39v4-Fc, demonstrated anti-viral effects against diverse Omicron subvariants, including BA4/5, BA.2.75, XBB, and BQ1, without the emergence of viral escape mutations. Compared to the intravenous administration of 3N39v4-Fc, the inhalation route required only twenty-fold lower dose to exhibit therapeutic effects in rodent models. Therapeutic potential of 3N39v4-Fc was also confirmed in a non-human primate model. Our accumulating evidence indicates that 3N39v4-Fc is the promising drug candidate for the treatment of SARS-CoV-2 infections even after complete escape from clinical antibodies.

## METHODS

### Plasmid generation

A plasmid encoding the SARS-CoV-2 Spike was obtained from Addgene #145032, and Omicron’s mutations and ΔC19 deletion (with 19 amino acids deleted from the C terminus) were cloned into pcDNA4TO (Thermo Fisher Scientific, Waltham. MA, USA)) (1). psPAX2-IN/HiBiT, pLenti Firefly, pCMV14 3N39v4, pCMV14 WT-ACE2 (1), and pcDNA3.4 Spike-foldon-his were described previously (42). The cDNA of WT or high-affinity ACE2 (3N39v4) was cloned into pCAGGS (43), together with three GGGS spacers and a 6xHis tag at the C-terminus. All cloning experiments were performed using an In-Fusion HD Cloning Kit (Takara Bio Inc., Shiga, Japan), and plasmid sequences were confirmed using a DNA sequencing service (Core Instrumentation Facility, Research Institute for Microbial Diseases, Osaka University, Osaka, Japan).

### Cell culture

Lenti-X 293T cells (Takara Bio Inc.) and its derivative, 293T/ACE2 cells, and Huh7/ACE2 (44) were cultured at 37 °C with 5% CO_2_ in Dulbecco’s modified Eagle’s medium (DMEM, WAKO) containing 10% fetal bovine serum (FBS) (Gibco) and penicillin/streptomycin (100 U/mL, Invitrogen) (P/S). Vero/TMPRSS2 cells (JCRB Cell Bank, Cat# JCRB1818) were cultured in DMEM containing 5% FBS and 1 mg/mL G418 (Roche, Basel, Switzerland). VeroE6/TMPRSS2 cells (JCRB Cell Bank, Cat# JCRB1819) were cultured in DMEM supplemented with 7% FBS and 1 mg/mL G418. All cell lines routinely tested negative for mycoplasma contamination.

### Viruses

SARS-CoV-2 (Wuhan:2019-nCoV/Japan/TY/WK-521/2020, Omicron (BA1.1: hCoV-19/Japan/TY38-871/2021, BA.1.18: hCoV-19/Japan/TY38-873/2021, BA.2: hCoV-19/Japan/TY40-385/2022, BA.2.75: hCoV-19/Japan/TY41-716/2022, XE: hCoV-19/Japan/TY41-686/2022, BA.4: hCoV-19/Japan/TY41-703/2022, BA.5: hCoV-19/Japan/TY41-702/2022) strain was isolated at National Institute of Infectious Diseases (NIID). The delta isolate (B.1.617.2 lineage; strain TKYTK1734) was provided by the Tokyo Metropolitan Institute of Public Health. These viruses were propagated in Vero/TMPRSS2 and VeroE6/TMPRSS2 cells. The tissue culture 50% infectious dose (TCID_50_) of the virus stocks were measured using VeroE6/TMPRSS2 cells.

Replication-competent vesicular stomatitis virus (VSV-SARS) was kindly provided by BEI Resources (NR-55284). VSV-SARS was propagated in Huh7/ACE2 cells. Mice adapted-SARS-CoV-2 (23) were generated using circular polymerase extension reaction (CPER), as described previously (44,45).

### Flow Cytometry

Expi293F cells (Thermo Fisher Scientific) were cultured at 37 °C, 130 rpm, 8% CO_2_ in Expi293 Expression Medium (Thermo Fisher Scientific). Cells were transfected with pcDNA4TO HMM-HA-Spike and pcDNA tagBFP for transfection marker using ExpiFectamine 293 (Thermo Fisher Scientific). Cells were centrifuged at 200 g for 2 min 48 h after transfection and washed with Dulbecco’s phosphate buffered saline (PBS) containing 0.2% bovine serum albumin (BSA). Cells were incubated for 30 min on ice with a serial dilution of ACE2-Fc in PBS-BSA. Cells were washed twice and resuspended for 30 minutes on ice in 1/250 anti-human IgG1-Fc-FITC (clone HP6017, BioLegend) and 1/1000 anti-hemagglutinin (HA) Alexa Fluor 647 (clone TANA2,1:4000 dilution; MBL). Cells were washed twice, resuspended in PBS-BSA, and analyzed on Attune NxT Flow Cytometer (Invitrogen). The main cell population was gated by forward-side scattering. Within the BFP-positive population with plasmid transfection, spike expression was analyzed with the mean fluorescence intensity of Alexa Fluor 647 and the binding of spike and ACE2-Fc was measured based on the mean FITC fluorescence intensity. EC_50_ were determined with the GraphPad Prism software.

### Protein synthesis and purification

Monoclonal antibodies, SARS-CoV-2 spike protein, and the engineered ACE2 were expressed using the Expi293F cell expression system (ThermoFisher Scientific) according to the manufacturer’s protocol. Fc-fused proteins were purified from the conditioned media using rProtein A Sepharose Fast Flow (Cytiva). Fractions containing target proteins were pooled and dialyzed against PBS.

The ACE2-6xHis or spike expression plasmid was transfected into Expi293F cells (A14635, ThermoFisher) using polyethyleneimine MAX (PEI MAX, 24765, Polyscience), and the cells were cultured in HE400AZ medium (Gibco). After five days, the cell culture supernatant was harvested and diluted to 20 mM Tris-HCl and 20 mM imidazole (pH 8.0) in the presence of a protease inhibitor cocktail (03969-34, Nacalai). The supernatant was loaded onto a Ni-NTA column pre-equilibrated with equilibration buffer (10 mM imidazole [pH 8.0], 20 mM Tri-HCl [pH 8.0], 0.3 M NaCl). The column was rinsed with washing buffer (20 mM imidazole [pH 8.0], 20 mM Tri-HCl [pH 8.0], 0.3 M NaCl) and eluted with elution buffer (250 mM imidazole [pH 8.0], 20 mM Tris-HCl [pH 8.0], 0.3 M NaCl). The eluted samples were purified using the Superose 6 Increase 10/300 GL column (Cytiva), equilibrated with 20 mM Tris-HCl (pH 8.0) and 250 mM NaCl buffer using an AKTA pure 25 chromatography system (Cytiva).

### VSV-SARS neutralization assay and virus passages

Huh7/ACE2 cells were seeded at 30,000 cells/well in 96-well plates. VSV-SARS was infected at an MOI of 0.1, together with ACE2-Fc protein or neutralizing antibody. Cellular expression of GFP, indicating viral infection, was determined using an imaging cytometer (BZX-800, Keyence). Viral supernatants with approximately 50% neutralization were used for subsequent viral passages. To determine the sequences of the S regions of VSV-SARS, the RNAs of the viral supernatants was isolated using ISOGENE II (NIPPON GENE). cDNA was synthesized using the PrimeScript™ II 1^st^ strand cDNA Synthesis Kit (Takara). PCR was performed using PrimeSTAR GXL Premix Fast, Dye Plus (Takara), and primers (46) (5′-CAGAGATCGATCTGTTTCCTTGACACGCGTGCCACCATGTTCGTGTTCCTG-3′ and 5′-AATCTGTGTGCAGGGCGGCCGCTCAGGTGTAGTGCAGCTTCACG-3′) and cloned by using the Zero Blunt TOPO PCR Cloning Kit (ThermoFisher).

### Pseudotyped virus neutralization assay

Pseudotyped reporter virus assays were conducted as previously described (1). Spike pseudovirus with a luciferase reporter gene was prepared by transfecting plasmids (pcDNA4 SARS-CoV-2 Spike, psPAX2-IN/HiBiT (47), and pLenti firefly) into LentiX-293T cells using Lipofectamine 3000 (Invitrogen). After 48 h, supernatants were harvested, filtered with a 0.45 μm low protein-binding filter (SFCA), and frozen at −80 °C. Then, 293T/ACE2 cells were seeded at 10,000 cells/well in a 96-well plate. HiBit value-matched pseudovirus and a three-fold dilution series of serum or therapeutic agents were incubated for 1 h, and this mixture was administered to ACE2/293T cells. After 1 h pre-incubation, the medium was changed, and cellular expression of the luciferase reporter indicating viral infection was determined using the ONE-Glo™ Luciferase Assay System (Promega) 48 h after infection. Luminescence was measured using an Infinite F200 Pro System (Tecan). Each serum sample was assayed in triplicate, and the 50% neutralization titer was calculated using Prism 9 (GraphPad Software).

### SARS-CoV-2 neutralization assay

Vero/TMPRSS2 cells were seeded at 80,000 cells in 24 well plates and incubated overnight. SARS-CoV-2 was then infected at an MOI of 0.1, together with 3N39v4. After 2 h, cells were washed with fresh medium and incubated with a fresh medium for 22 h. The culture supernatants were collected and subjected to qRT-PCR.

### Rodent models of SARS-CoV-2 infection

Four-week-old male Syrian hamsters were purchased from SLC Japan. Syrian hamsters were anaesthetized by isoflurane and challenged with 1.0 × 10^6^ TCID_50_ (in 60 μL) via intranasal routes. After 2 h post infection, 3N39v4 (20 mg/kg) was administered through intraperitoneal injection. Five days post-infection, all animals were euthanized, and the lungs were collected for virus titration and qRT-PCR.

In the BALB/c mouse model, 10-week-old female BALB/c mice were purchased from SLC Japan or CLEA Japan. Mice were anaesthetized by isoflurane and challenged with MA10 (1.0 × 10^4^ TCID_50_ in 20 μL) via intranasal routes. Then, 24 h after infection, 3N39v4 (0.25-20 mg/kg) was administered through intraperitoneal injection or inhalational administration as described above. Five days post-infection, the animals were euthanized, and lungs were collected for qRT-PCR and virus titration.

All animal experiments on SARS-CoV-2 were performed in biosafety level 3 (ABSL3) facilities at the Research Institute for Microbial Diseases, Osaka University (Osaka, Japan). Animal experimentation protocols were approved by the Institutional Committee of Laboratory Animal Experimentation of the Research Institute for Microbial Diseases, Osaka University (approval number R02-08-0). All efforts were made during the study to minimize animal suffering and reduce the number of animals used in the experiments.

### Monkey model of SARS-CoV-2 infection

Nine adult male cynomolgus monkeys (CMs) (4-8 years of age, 2.9-4.2 kg weight) housed at the Tsukuba Primate Research Center (TPRC), National Institutes of Biomedical Innovation, Health, and Nutrition (NIBIOHN, Ibaraki, Japan) were used in this study. All animals were negative for the B virus, simian immunodeficiency virus, simian T-cell leukemia virus, and *Mycobacterium*. The animals were handled under the supervision of the veterinarians in charge of the animal facility.

The monkey experiments were approved by the Committee on the Ethics of Animal Experiments of NIBIOHN (approval number: DSR03-36) under the guidelines for animal experiments at NIBIOHN and in accordance with the Guidelines for Proper Conduct of Animal Experiments established by the Science Council of Japan (http://www.scj.go.jp/ja/info/kohyo/pdf/kohyo-20-k16-2e.pdf). The experiments were conducted in accordance with the “Weatherall report for the use of non-human primates in research” recommendations (https://royalsociety.org/topics-policy/publications/2006/weatherall-report/). Animals were housed in adjoining individual primate cages, allowing them to make sight and sound contact with one another for social interactions; temperature was maintained at 25°C, with light for 12 h per day. The animals were fed apples and a commercial monkey diet (Type CMK-2; Clea Japan, Inc., Tokyo, Japan).

Infection experiments were performed as previously described (48). Briefly, the animals were housed in ABSL3 facilities at the TPRC of NIBIOHN during the experimental period and monitored throughout the study for physical health and clinical assessment. The clinical status of CMs was scored daily in six categories as follows: appearance (skin and fur; 0–10), secretion (nose, mouth, eyes; 0–5), respiration (0–15), discharge (feces and urine; 0–10), appetite (food intake; 0–10), and activity (0–15) (49). A data logger (DST milli-HRT; Star-Oddi) was embedded intraperitoneally into all CMs to monitor the body temperature every four hours daily. CMs were inoculated with SARS-CoV-2 Delta isolate under anesthesia in a BSL3 room via a combination of i.t. (900 μL) and i.n. (50 μL per nostril) routes, with a total of 1 × 10^6^ TCID_50_ virus. All animal experiments were conducted under anesthesia to obtain blood and swab samples and for CT scanning at each collection point. The animals were euthanized at the end of the experiment. At euthanasia, the animals were deeply anesthetized with pentobarbital under ketamine anesthesia, and whole blood samples were collected from the left ventricle.

### Micro-CT imaging of infected CMs

Micro-CT imaging of the infected macaques was performed as previously described (48). Briefly, under anesthesia (7.5 mg/kg ketamine⋅HCl and 3 mg/kg xylazine), CT images were obtained (80 kV, 400 μA; field of view,160 mm; scan time, respiratory gated, 8 min). All CT scans were performed at a BSL3 facility using a 3D micro-CT scanner system (Cosmo Scan CT AX; Rigaku Corporation, Tokyo, Japan). Lung images were reconstructed using the CosmoScan Database software of the micro-CT scanner (Rigaku Corporation). Slices of the third, sixth, and ninth thoracic vertebrae, including the upper, middle, and lower lung areas, were selected. Images were analyzed using a Cosmo Scan CT viewer (Rigaku Corporation).

### Histological analysis

From each lung, the left lower, right lower, left upper, and right upper lobes were cut and processed routinely to prepare formalin-fixed paraffin-embedded tissue samples. The samples were cut into 2 μm-thick tissue sections and stained with H&E. H&E-stained tissue sections were scanned using SLIDEVIEW VS200 (Olympus Life Science, Tokyo, Japan) to acquire whole-slide digital virtual slide images. Using the image analysis software QuPath, the algorithm to classify the histological pattern of the inflammatory area was trained, and the ratio of the inflammatory area to the total lung tissue area was calculated.

### Quantitative RT-PCR of *in vivo* samples

In small animal experiments, total RNA from lung homogenates was isolated using ISOGENE II (NIPPON GENE, Tokyo, Japan)). Real-time RT-PCR was performed using the Power SYBR Green RNA-to-CT 1-Step Kit (ThermoFisher Scientific) on the AriaMx Real-Time PCR system (Agilent, Santa Clara, CA, USA)). Relative quantitation of target mRNA levels was performed using the 2-ΔΔCT method. The values were normalized to those of the housekeeping gene, β-actin. The following primers were used: β-actin (hamster/mouse), 5′-TTGCTGACAGGATGCAGAAG-3′ and 5’-GTACTTGCGCTCAGGAGGAG-3; 2019-nCoV_N2, 5′-AAATTTTGGGGACCAGGAAC -3′and 5′-TGGCAGCTGTGTAGGTCAAC -3′; IL-6 (hamster), 5′-GGA CAATGACTATGTGTTGTTAGAA -3′ and 5′-AGGCAAATTTCCCAATTGTATCCAG -3′; CCL3 (hamster), 5′-GGTCCAAGAGTACGTCGCTG -3′ and 5′-GAGTTGTGGAGGTGGCAAGG -3′; CCL5 (hamster), 5′-TCAGCTTGGTTTGGGAGCAA -3′ and 5′-TGAAGTGCTGGTTTCTTGGGT -3′; CXCL10 (hamster), 5′-TACGTCGGCCTATGGCTACT -3′ and 5’-TTGGGGACTCTTGTCACTGG -3′; and IL-6 (mouse); 5′-CCACTTCACAAGTCGGAGGCTTA -3′ and 5′-GCAAGTGCATCATCGTTGTTCATAC -3′.

In monkey experiments, quantification of viral RNA in swab samples was performed as described previously (48). Briefly, nasal and pharyngeal swab samples were collected in 1 mL PBS supplemented with an Antibiotic-Antimycotic Mixed Stock Solution (Nacalai Tesque, Kyoto, Japan). RNA was extracted from swab samples using a QiaAmp Viral RNA kit (Qiagen), according to the manufacturer’s instructions. To detect viral RNA, 5 μL of extracted RNA was subjected to one-step real-time RT-PCR using a One Step PrimeScript™ RT-PCR Kit (Perfect Real Time) (TaKaRa Bio Inc.) on a QuantStudio 5 Real-Time PCR system (ThermoFisher Scientific). We used Primer/Probe N2 (2019-nCoV) (TaKaRa Bio Inc.). The reaction conditions for RT-PCR were 42 °C for 5 min and 95 °C for 10 s, followed by 45 cycles of 5 s at 95 °C for 34 s and 60 °C.

To examine the viral distribution in infected CMs, viral RNA levels in the tissue samples were quantified at necropsy, as described previously (48). Briefly, samples were collected from the same location and the same position of each lung lobe, tissue, and organ using a biopsy-punch instrument (5 mm; Kaijirushi). All samples collected from the tissues and organs were placed in a DNA/RNA Shield (Zymo Research, CA, USA). Tissue samples were homogenized using gentleMACS M Tubes (Miltenyi Biotec, Bergisch Gladbach, Germany) with a gentleMACS Dissociator (Miltenyi Biotec), and RNA was extracted using an RNeasy Mini Kit (Qiagen, Venlo, Netherland). The extracted RNAs were subjected to one-step real-time PCR using the One Step PrimeScript™ RT-PCR Kit (Perfect Real Time) (TaKaRa Bio Inc.).

To quantify the copy number of the N gene, we used Primer/Probe N2 (2019-nCoV) (TaKaRa). The reaction conditions for RT-PCR were 42 °C for 5 min and 95 °C for 10 s, followed by 45 cycles of 5 s at 95 °C for 34 s and 60 °C. For quantification of subgenomic RNA (sgRNA), we used the forward primer 5′-CGATCTCTTGTAGATCTGTTCTC-3′ and reverse primer 5′-ATATTGCAGCAGTACGCACACA-3′ with the probe FAM-5′-ACACTAGCCATCCTTACTGCGCTTCG-3′-BHQ1. The reaction conditions were 42 °C for 5 min and 95 °C for 10 s, followed by 45 cycles of 5 s at 95 °C for 34 s and 60 °C.

### Radiolabeling and PET analysis

Coupling of the bifunctional chelate and radiolabeling was performed as previously described by Vosjan et al (50). Briefly, a three-fold equivalent of DFO-Bz-NCS in DMSO (less than 2% of the total volume) was added to the ACE2 Fc protein (1–2 mg) dissolved in PBS adjusted to pH 8.9-9.1. The reaction mixture was incubated in a water bath at 37 °C for 1 h. Purification was performed using a size-exclusion chromatography column (PD-10; GE Healthcare, Chicago, IL, USA) and eluted with PBS. The neutralized 89Zr-oxalate solution (30-50 MBq) with 2 M sodium acetate was mixed with 0.6-0.7 mL of 0.5 M HEPES, 80-140 μL of gentisic acid in 0.25 M sodium acetate (pH 5.4–5.6) and the protein solution. The reaction mixture was incubated at room temperature for 1 h. Free 89Zr metal was separated using a size exclusion chromatography column (GE Healthcare) and ^89^Zr-DFO-3N39v4-Fc. The radiochemical purity was estimated to be > 95% based on TLC autoradiography (TLC-ARG) and HPLC (LC-20; Shimadzu) using a Superdex 200 10/300 column (GE Healthcare). ^89^Zr-DFO-3N39v4-Fc was administered into the tail vein (400ug, 0.84-1.0MBq) or inhalation routes (80ug, 1.75-2.0 MBq) to C57BL/6j mice (n = 5), and PET data was obtained at 0-7 days after administration using a PET system (Clairvivo PET; Shimadzu.JAPAN). CT data were obtained using a PET/CT system (Eminence Stargate; Shimadzu. JAPAN). The acquired PET data were reconstructed using 3-D-DRAMA. PET and CT images were converted to the DICOM format and fused using PMOD software version 3.3 (PMOD Technologies Ltd.). Regions of interest were drawn from axial images in fused PET/CT images. The standard uptake value (SUV) was calculated as follows: SUV = [tissue radioactivity in Bq/g tissue]/[injected radioactivity in Bq/g body weight].

### Inhalational administration

BALB/c mice or Syrian hamsters were anaesthetized by the intraperitoneal administration with 10 μL/g body weight of a medetomidine-midazolam-butorphanol tartrate mixture (0.75 mg/kg-medetomidine (ZENOAQ), 2 mg/kg midazolam (Maruishi-pharm), and 2.5 mg/kg butorphanol tartrate (Meiji Seika). ACE2-FC in 50 μL PBS for mice or 200 μL PBS for hamster was administered through inhalation using an aerosol sprayer (Natsume Seisakusho, KN-34700-1). Atipamezole (ZENOAQ) was administered intraperitoneally to recover from anesthesia.

### Pharmacokinetics of ACE2-Fc in mice by inhalation administration

Briefly, 0.25 mg/kg or 1.0 mg/kg 3N39v4 were administered through inhalation as described above. Mice were euthanized, and lung tissues were collected at different time points (0, 2, 6, 12, 24, 48, and 72 h). Lung tissues were weighed and homogenized in lysis buffer, consisting of 20 mM Tris-HCl (pH 7.4), 135 mM NaCl, 1% Triton X-100, 1% glycerol, and protease inhibitor cocktail (Roche Molecular Biochemicals), using gentleMACS M Tubes (Miltenyi Biotec) with a gentleMACS Dissociator (Miltenyi Biotec), and centrifuged at 13,000 rpm. The supernatant was harvested and stored at −80 °C for further quantification. For immunoblotting, the supernatants were incubated with sample buffer at 95 °C for 5 min. Samples were resolved using SDS-PAGE (NuPAGE gel, Life Technologies) and transferred onto nitrocellulose membranes (iBlot, Life Technologies). The membranes were blocked with PBS containing 5% skim milk and incubated with HRP-conjugated anti-human IgG1 antibodies (Promega, #W403B) overnight at 4 °C.

The immune complexes were visualized with SuperSignal West Femto substrate (Pierce) and detected using an Image Quant 800 image analyzer system (Amersham). The 3N39v4 concentration was analyzed using an in-house sandwich ELISA, as described previously (5). In brief, 10 μg/mL goat anti-human IgG1 Fc F(ab’)2 (Sigma-Aldrich, I3391) was coated in 96-well NUNC Immunoplate (Thermo Fisher, #439454) overnight at 4 °C and and blocked with Blocking One (Nacalai Tesque) for 2 h at 25°C. Serially diluted lung homogenates were added to the plates and incubated for 2 hours at 37 °C. The plate was washed with TBS thrice and incubated with HRP-labeled anti-ACE2 IgG diluted at 0.4 µg/mL for 1 h at room temperature. After washing three times, the enzyme activity was measured using a GloMax Discover Microplate Reader (Promega), and absorbance was recorded at 405 nm after incubation with the ABTS substrate (Nacalai Tesque) for 10 min. ACE2-FC concentration was measured using standard curves obtained using serially diluted ACE2-FC in the same experiment.

### Cryo-EM imaging

To prepare the cryo-EM grids for WT-ACE2-spike or 3N39v4-spike complexes, 2.2 μL aliquots of the sample solution were applied to glow-discharged holey Quantifoil R0.6/1.0, Au 200 mesh grids (Quantifoil Micro Tools GmbH, Germany). The grids were blotted with filter paper for 6 s and immediately plunged into liquid ethane using a Vitrobot Mark IV (ThermoFisher Scientific) at 4 °C and 95% humidity. The frozen grids were subsequently transferred to a cryo-electron microscope (Titan Krios, ThermoFisher Scientific), equipped with a field emission gun and Cs corrector (CEOS, GmbH). The microscope was operated at 300 keV in nanoprobe and EF-TEM modes at the Institute for Protein Research, Osaka University. Images were acquired as movies using a Gatan BioQuantum energy filter (slit width of 20 eV) and K3 camera (Gatan, Inc., USA) in the electron counting mode. The 4,545 (WT) and 7,947 (3N39v4) images were collected at a nominal magnification of 81,000×, resulting in a final pixel size of 0.880 Å, and a total dose of 60 e-/Å2 was fractionated into 82 movie frames. SerialEM software (51) was used for automated data collection using a 3 × 3 pattern beam-image shift scheme with a nominal defocus range of −0.8 to −1.8 mm.

### Image processing

Image processing steps for each complex are summarized in Supplemental Figure 3 and S4. All steps were performed using the cryoSPARC v3.3.1 software (52). For the WT-ACE2 complex and the 3N39v4 complex 4,517, and 7,795 movies were used, respectively. After motion correction and contrast transfer function estimation, auto-picking on Topaz machine-learning (53) resulted in 793,794, and 880,910 particles for WT-ACE2 and the 3N39v4, respectively. The extracted particles were subjected to heterogenous refinement into eight classes using multi reference maps to distinguish the state of RBD and ACE2 binding. The reference maps were generated from PDB models (PDBID: 7DK3, 7DF4, 7KMZ, and 7KMS) using USCF chimera molmap command (54). Then, the particles belonging to good classes were subjected to *ab initio* reconstruction to remove junk particles by reconstructing three classes. The selected particles were subjected to multi-reference heterogenous refinement. Finally, particles of each class were homogenous refinement to obtain a high-resolution map. To obtain the detailed structure of the 3N39v4 and RBD, local refinement on this region was carried out after homogenous refinement. The density map for the 3N39v4 and RBD was obtained at 3.40 Å. The resolution was based on the gold standard criterium: Fourier shell correlation = 0.142 criterion.

### Model building and refinement

To build the atomic model of the 3N39v4 and RBD complex, that of the 3N39 and RBD complex (PDBID:7DMU) was used as an initial model. The initial model was fitted into the density map as a rigid body and flexibly fitted against the map using the Chain Refine function of COOT software (55). The roughly fixed model was extensively manually corrected, residue by residue, using the real-space refine zone function in COOT. The manually corrected model was refined using the phenix_real_space_refine program (56), with a secondary structure and Ramachandran restraints. The resulting model was manually checked using COOT. The water molecules were manually placed. This iterative process was performed for several rounds to correct the remaining errors until the model satisfied the geometry requirements. Model quality was assessed using Molplobity and EMRinger scores (57,58).

For model validation against overfitting, the final models were used to calculate the Fourier shell correlation against the final density maps used in model building using the phenix refine program. The statistics of the maps and models are summarized in Supplemental Table 1. All structures were prepared using the UCSF chimeraX software (59).

### Field potential recordings

FPs were recorded as previously described (60). Briefly, before measuring FPs, hiPS-CM were incubated for more than 30 min in a 5% CO_2_ incubator at 37 °C, after replacement with a fresh medium. To stabilize the waveform, the probes were set on an MEA system (MED64, Alpha MED Scientific) and equilibrated for at least 30 min in a humidified 5% CO_2_ atmosphere. After treatment with 3N39v4 (1%), FP recordings were performed for 10 min. Vehicle- and baseline-corrected field potential durations (ΔΔFPDc) were calculated for 3N39v4 (61). Dofetilide (3 nM), which causes prolongation of field potential duration and subsequent early afterdepolarization at 24 h, was used as a positive control.

### Motion analysis

The motion of hiPSC-CMs was recorded using the SI8000 cell motion imaging system (Sony, Japan), as previously reported (62). Sequential phase-contrast images were obtained using a 10× objective at a frame rate of 150 frames per sec and resolution of 2048 × 2048 pixels at a rate of 150 Hz for 10 s at 37 °C using the SI8000 cell motion imaging system (Sony, Japan). The recorded movie images were analyzed with a block-matching algorithm using an SI8000 analyzer (63).

### Membrane protein array

Approximately 6,000 native human membrane proteins were tested for interactions with 3N39v4 using a membrane protein array™ (Integral Molecular).

### T cell proliferation assay

The T cell proliferation assay was performed by Proimmune Inc. The peptides used in this study are listed in Supplemental Tables 2.

### Statistical analysis and illustration

Neutralization measurements were performed in technical triplicates, and relative luciferase units were converted to percent neutralization and plotted using a non-linear regression model to determine the IC50/ID50 values using the GraphPad PRISM software (version 9.0.0). Comparisons between two groups of paired data were performed using a paired *t* test. Comparisons between two groups were made with unpaired *t* tests, and log-rank tests. More than two groups were compared by one-way analysis of variance (ANOVA) followed by Dunnett’s multiple comparison test. P value < 0.05 was considered as statistically significant.

## Acknowledgments

We would like to thank all members of “Team Handai project meeting” for helpful discussion, Kenzo Tokunaga (Department of Pathology, National Institute of Infectious Diseases) for the kind gift of the plasmid coding psPAX2-IN/HiBiT, This work was supported by the Japan Agency for Medical Research and Development (AMED), Research Program on Emerging and Re-emerging Infectious Diseases under JP21fk0108465 (A.H., J.T., Y.Y., A.S. and Toru.O.), JP21fk0108481 (A.H., J.T., Y.Y., A.S., and Toru. O.) and JP22fk0108144 (Toru.O.), AMED Japan Program for Infectious Diseases Research and Infrastructure JP21wm0325009, and JP22wm0325009 (A.S. and Toru.O.), Platform Project for Supporting Drug Discovery and Life Science Research (Basis for Supporting Innovative Drug Discovery and Life Science Research) under JP21am0101075 (J.T.), Mochida Memorial Foundation for Medical and Pharmaceutical Research (Toru.O.), Grant for Joint Research Projects of the Research Institute for Microbial Diseases, Osaka University (A.S.), and MSD Life Science Foundation, Public Interest Incorporated Foundation (Y.H.).

## Author contribution

A.S. and Toru O. designed the study; E.U., Tomotaka.O., Shuya.M., Y.Y., A.S., and Toru.O. performed the cynomolgus monkey experiments; J.I.K., K.A., H.A., M.O. and T.K., and performed the CryoEM experiments; Y.I., T.S., A.T., Tadahiro.S., R.K., S.H., Tatsuo.S., E.H., and Toru.O. performed the SARS-CoV-2 cell-culture experiments and rodent models; M.K. and Toru.O. performed the VSV-SARS experiments; Takahiro.S. performed the PET analysis; Y.H., Satoaki.M., and A.H. performed the peudovirus experiments; Y.M. generated mouse-adapted SARS-CoV-2; J.T. purified and prepared the proteins; S.Y. and Y.K. evaluated cardiotoxicity; A.H., T.K., M.O., Y.Y., A.K., and Toru.O. supervised the research; E.U., Y.I., T.S., J.I.K., K.A., Y.H., A.H., Y.S., A.S., and Toru. O. wrote the manuscript. All authors discussed the results and commented on the manuscript.

