## Supplementary data for "An engineered ACE2 decoy broadly neutralizes Omicron subvariants and shows therapeutic effect in SARS-CoV-2-infected cynomolgus macaques"

**A**

| variant | mutations | Ref. |
|---|---|---|
| 3N39v2 | A25V, K31N, E35K, L79F S128C, V343C | 4 |
| 3N39v3 | A25V, K31N, E35K S128C, V343C | |
| 3N39v4 | A25V, K31N, E35K, T92Q S128C, V343C | |
| 3J113v2 | K31M, E35K, Q60R, L79F | 4 |
| 3J320v2 | T20I, H34A, T92Q, Q101H | 4 |

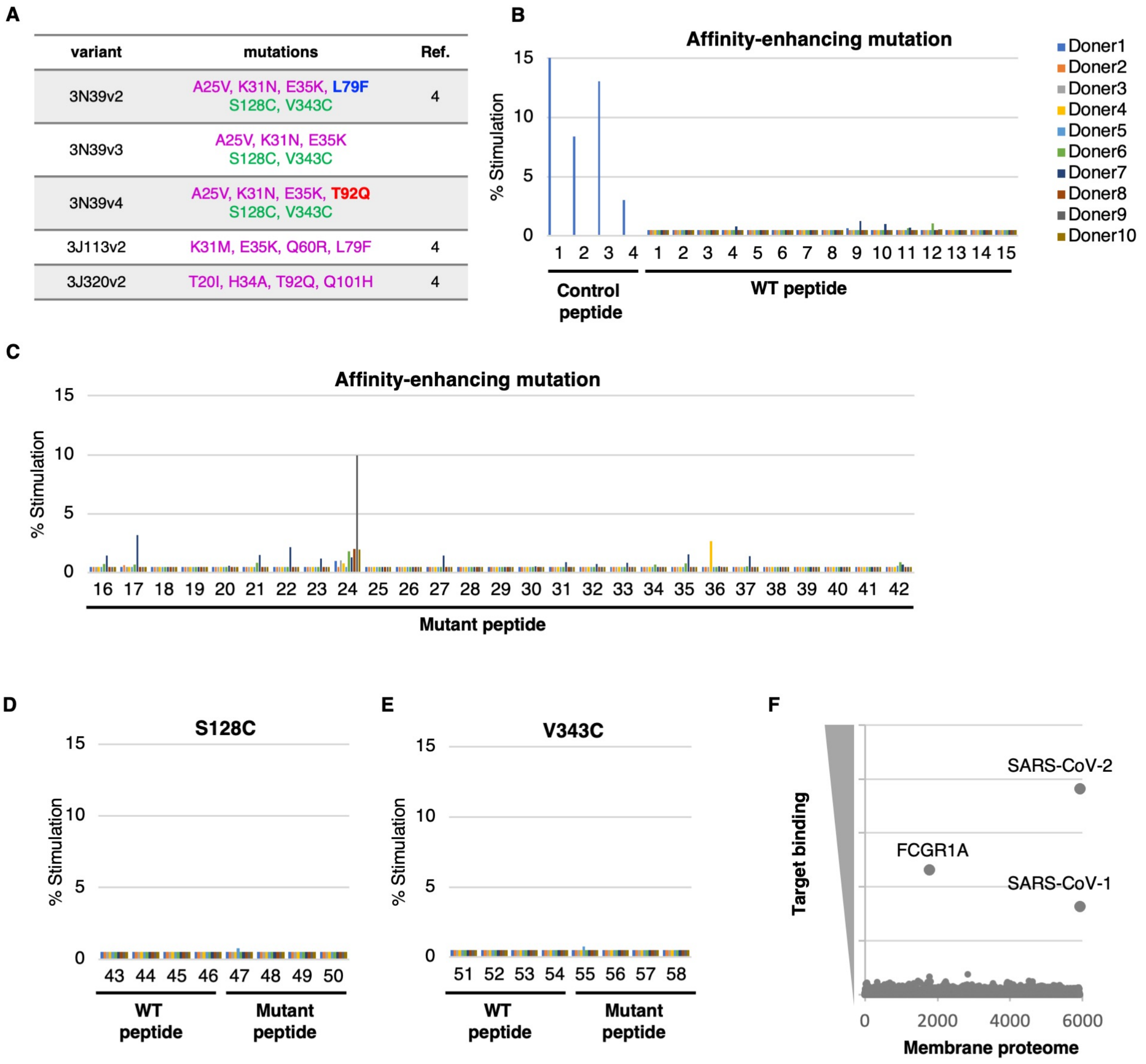


**Supplemental Figure 1. Screening of antigenicity for high affinity ACE2 decoys.**

**(A)** Amino acid substitutions are listed for affinity enhancement and peptidase dead in each ACE2 mutants. **(B, C)** Peripheral blood mononuclear cells (PBMC) derived from 10 different HLA donors were incubated with control peptides and wild-type (WT) ACE2 peptides (B) and affinity-enhancing mutant peptides (C), as described in Tables S1. Peptide #24 contains F79 stimulated T cells from multiple donors. **(D, E)** PBMCs derived from 10 different HLA donors were incubated with WT and mutant peptides described in Tables S1. Antigenicity of S128C (D) and V343C (E) was independently evaluated with different 10 HLA donor set. **(F)** Flow cytometry-based membrane proteome array was conducted to identify endogenous off-target binding proteins of 3N39v4-Fc**.**

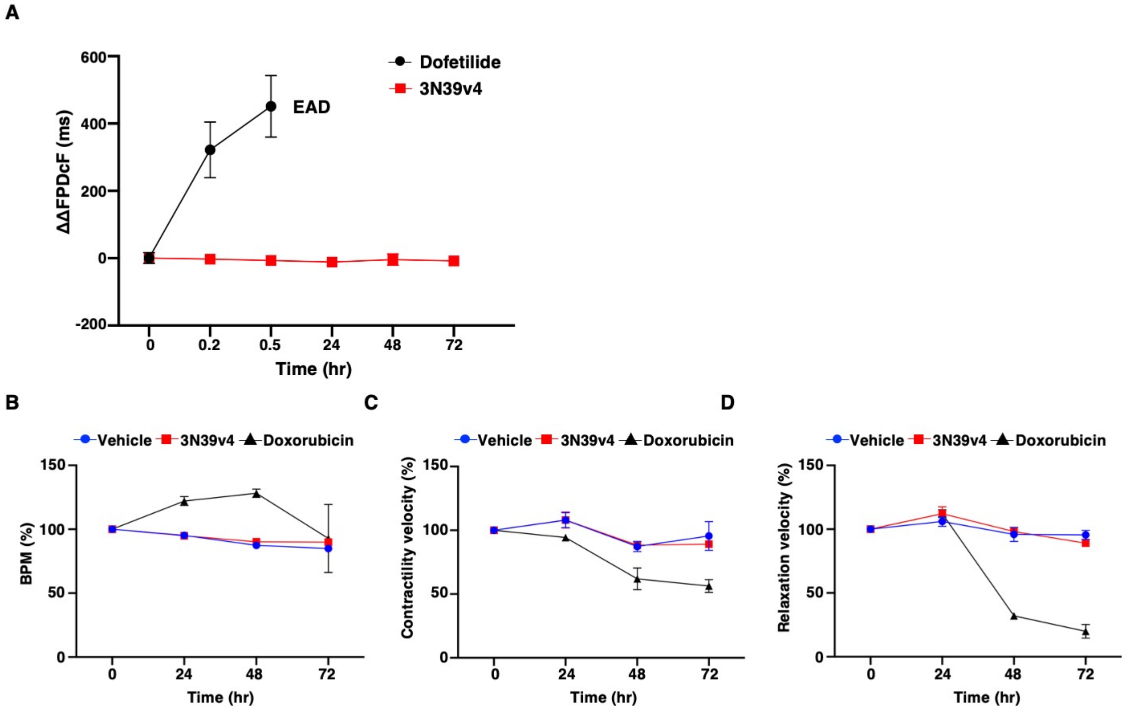


**Supplemental Figure 2. Evaluation of cardiotoxicity using human iPS-derived cardiomyocytes (hiPS-CMs). (A)** Change in ΔΔFPDcF following treatment with 3N39v4 (100 μg/mL) and positive control, dofetilide (3 nM), were measured by Field potential recordings. **(B-D)** The effect of 3N39v4 (100 μg/mL) and positive control, doxorubicin (1 μM), on the contraction parameters were measured by motion analysis (B, BPM; C, contraction velocity; D, relaxation velocity).

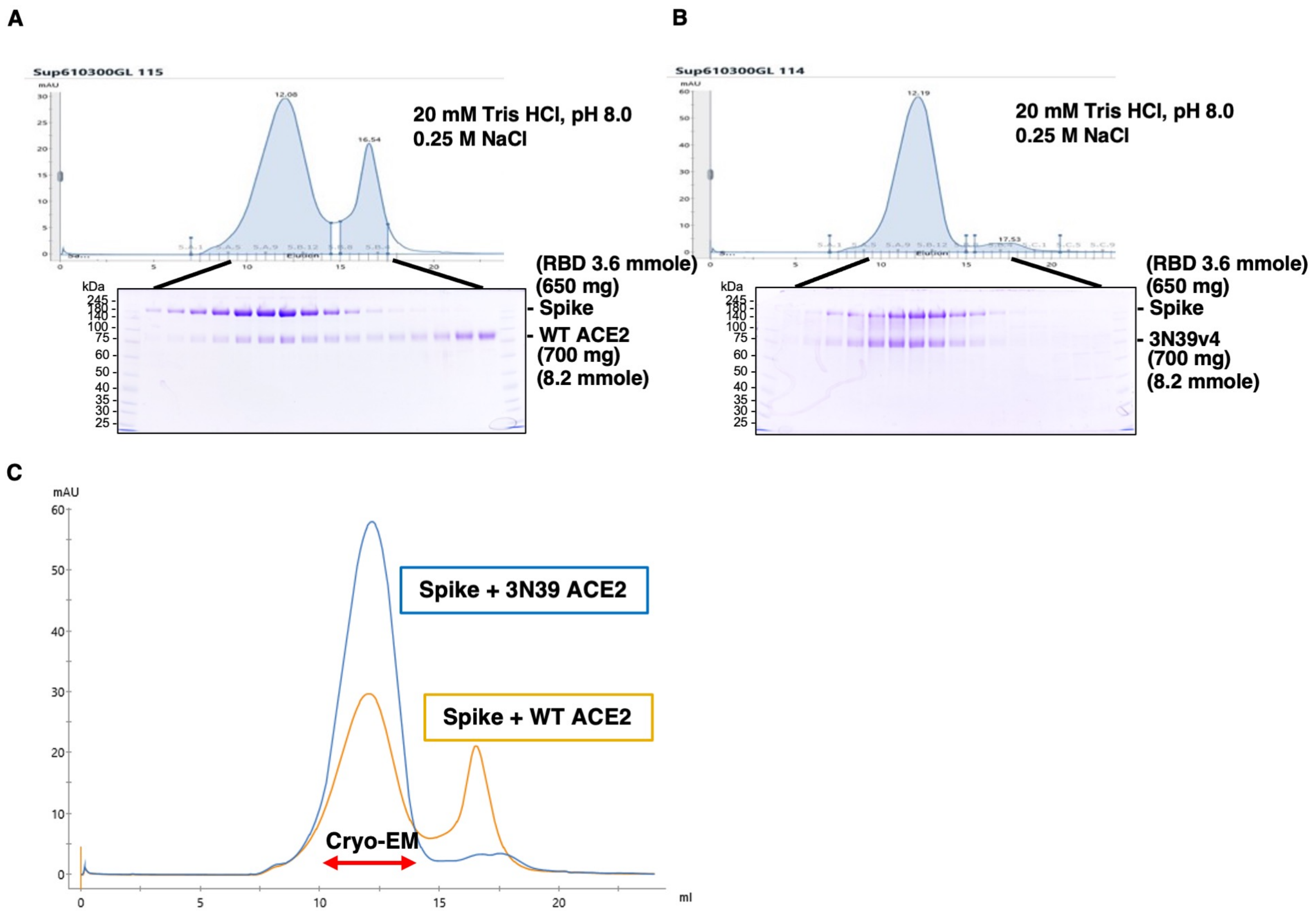


**Supplemental Figure 3. Biochemical analysis of the interaction between spike protein and wild-type ACE2 or 3N39v4.**

**(A)** Gel filtration chromatography of spike protein mixed with an excess amount of WT ACE2. **(B)** Gel filtration chromatography of spike protein mixed with an excess amount of 3N39v4. **(C)** Merged image of two chromatograms of wild-type ACE2 and SARS-CoV-2 spike proteins (yellow, panel A) and 3N39v4-ACE2 and SARS-CoV-2 spike proteins (blue, panel B). The purified fractions indicated by the red arrow were analyzed using Cryo-EM.

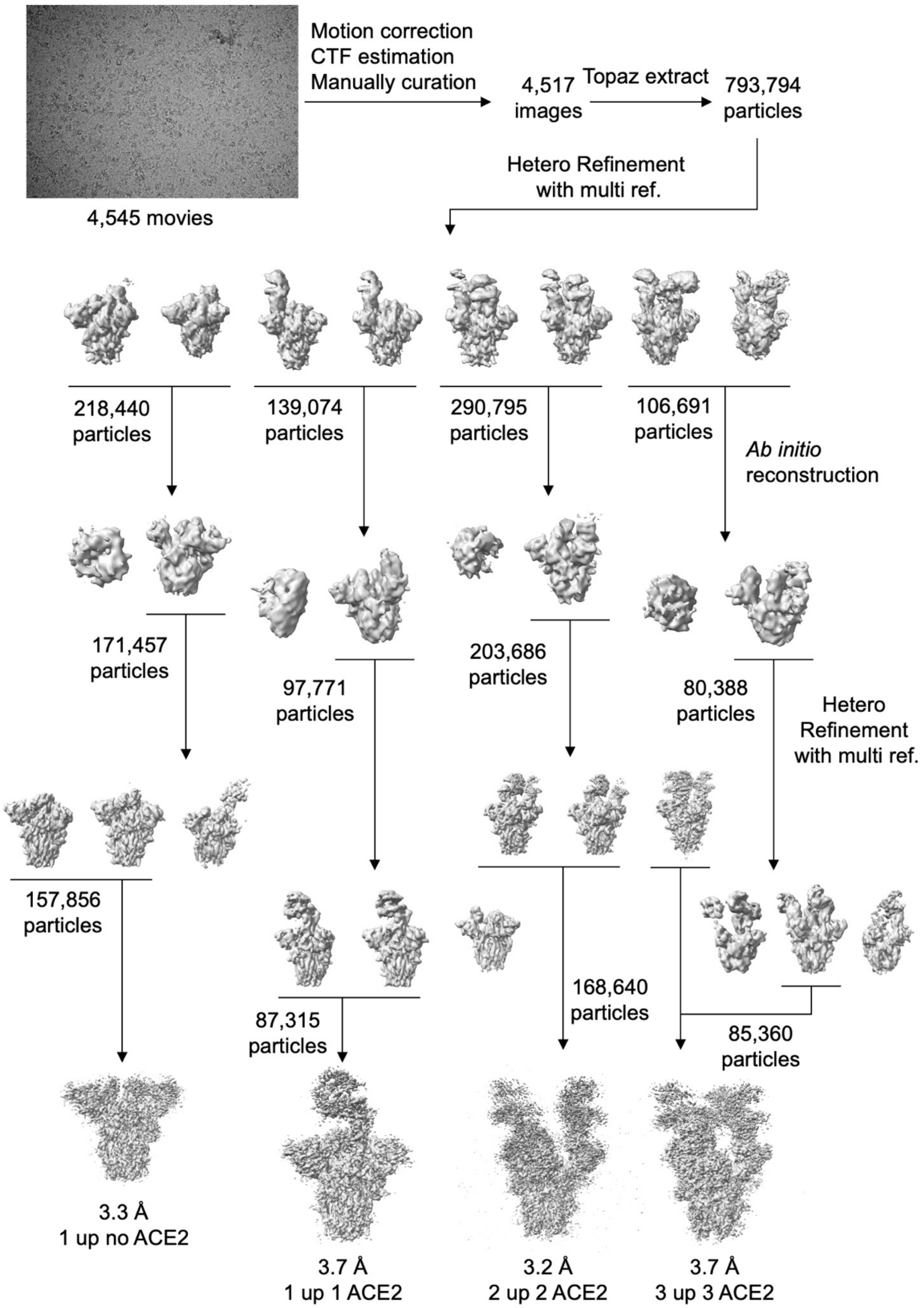


**Supplementary Figure 4. Flow chart of single particle cryoEM for the complex of spike protein and WT-ACE2.**

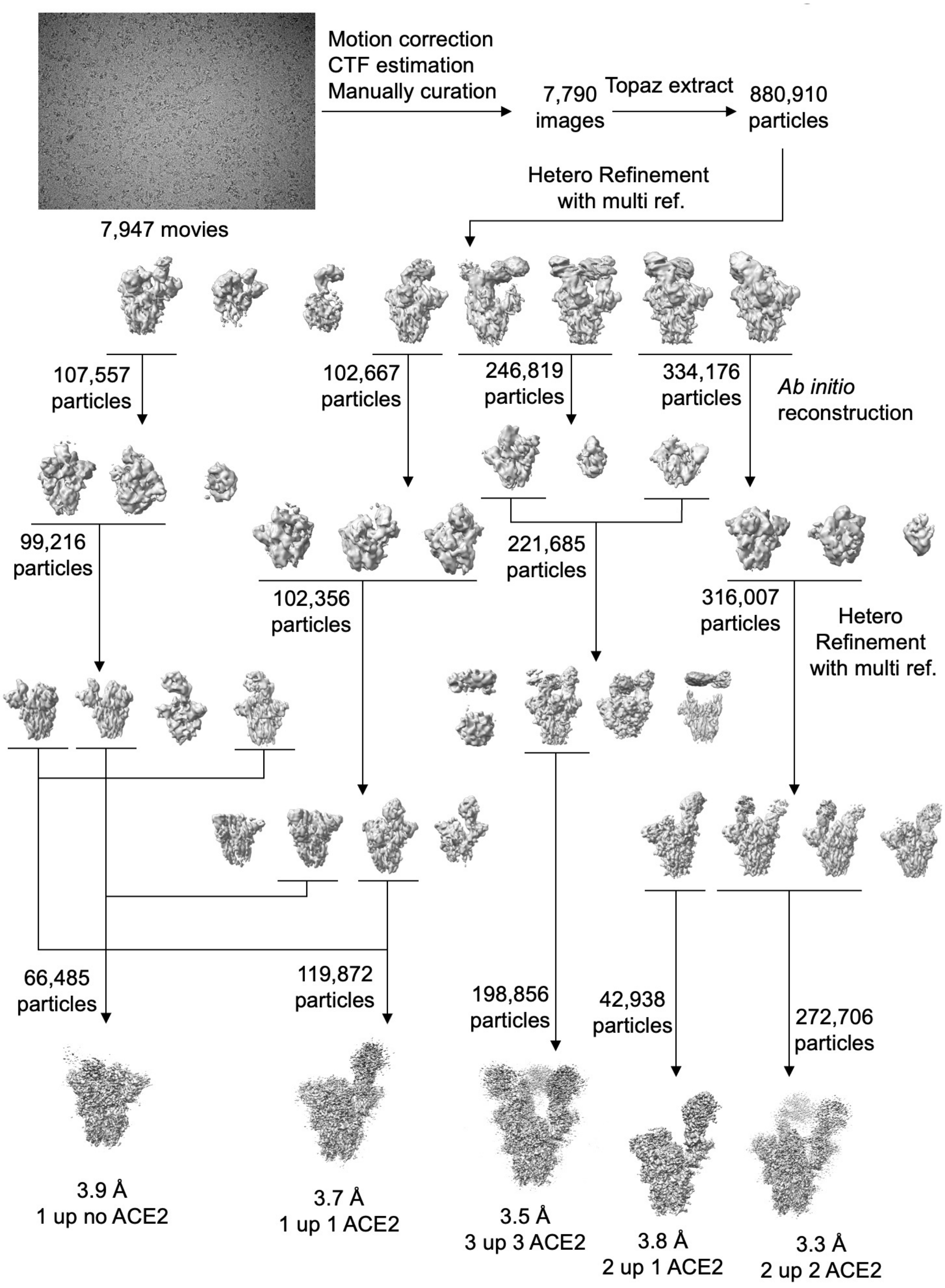


**Supplementary Figure 5. Flow chart of single particle cryoEM for the complex of spike protein and ACE2 decoy (3N39v4).**

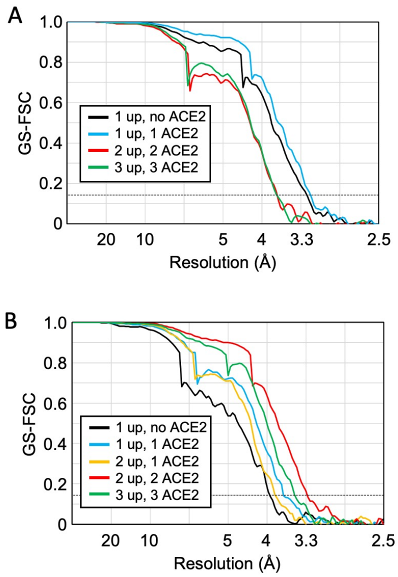


**Supplemntary Figure 6. GS-FSC curves for each condition.** (A) The complex of spike protein and WT-ACE2. (B) The complex of spike protein and ACE2 decoy (3N39v4). Black dotted line shows GS-FSC=0.143 criteria.

A

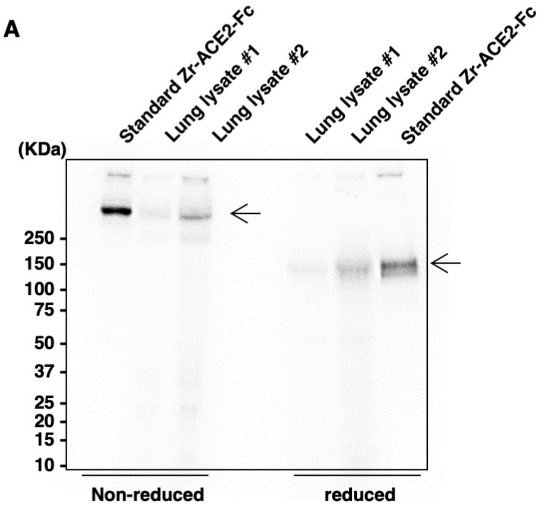


B

| | | Non-Reduced | | Reduced | |
|---|---|---|---|---|---|
| | | STD | Lung lysate | STD | Lung lysate |
| Day1 | Total (PSL) | 26897.1 | 18988.8 | 29792.1 | 19497.9 |
| | ACE2 (PSL) | 21359.3 | 5804.4 | 26326.0 | 10813.3 |
| | Ratio (%) | 79.4 | 30.1 | 88.4 | 54.3 |
| Day4 | Total (PSL) | 12170.8 | 7555.8 | 14788.6 | 6254.7 |
| | ACE2 (PSL) | 9993.6 | 1862.8 | 13196.9 | 2665.2 |
| | Ratio (%) | 82.1 | 24.6 | 89.2 | 42.5 |
| Day7 | Total (PSL) | 7493.3 | 3755.5 | 7821.2 | 3712.2 |
| | ACE2 (PSL) | 5905.3 | 534.7 | 6797.5 | 1132.6 |
| | Ratio (%) | 78.8 | 14.2 | 86.9 | 29.7 |

**Supplemental Figure 7. Pharmacokinetics of inhaled 3N39v4-Fc.**

**(A)** Imaging of SDS-PAGE for lung lysate in non-reduced and reduced condition 24 h after inhalation of $^{89}$Zr-labelled 3N39v4-Fc at a dose of 80 µg/body. Arrow indicates dimer or monomer of 3N39v4-Fc in non-reduced or reduced condition, respectively. **(B)** The value of photo-stimulated luminescence (PSL) of pre-loading total lysate and ACE2 segment after electrophoresis and the ratio of ACE2 value per total value.

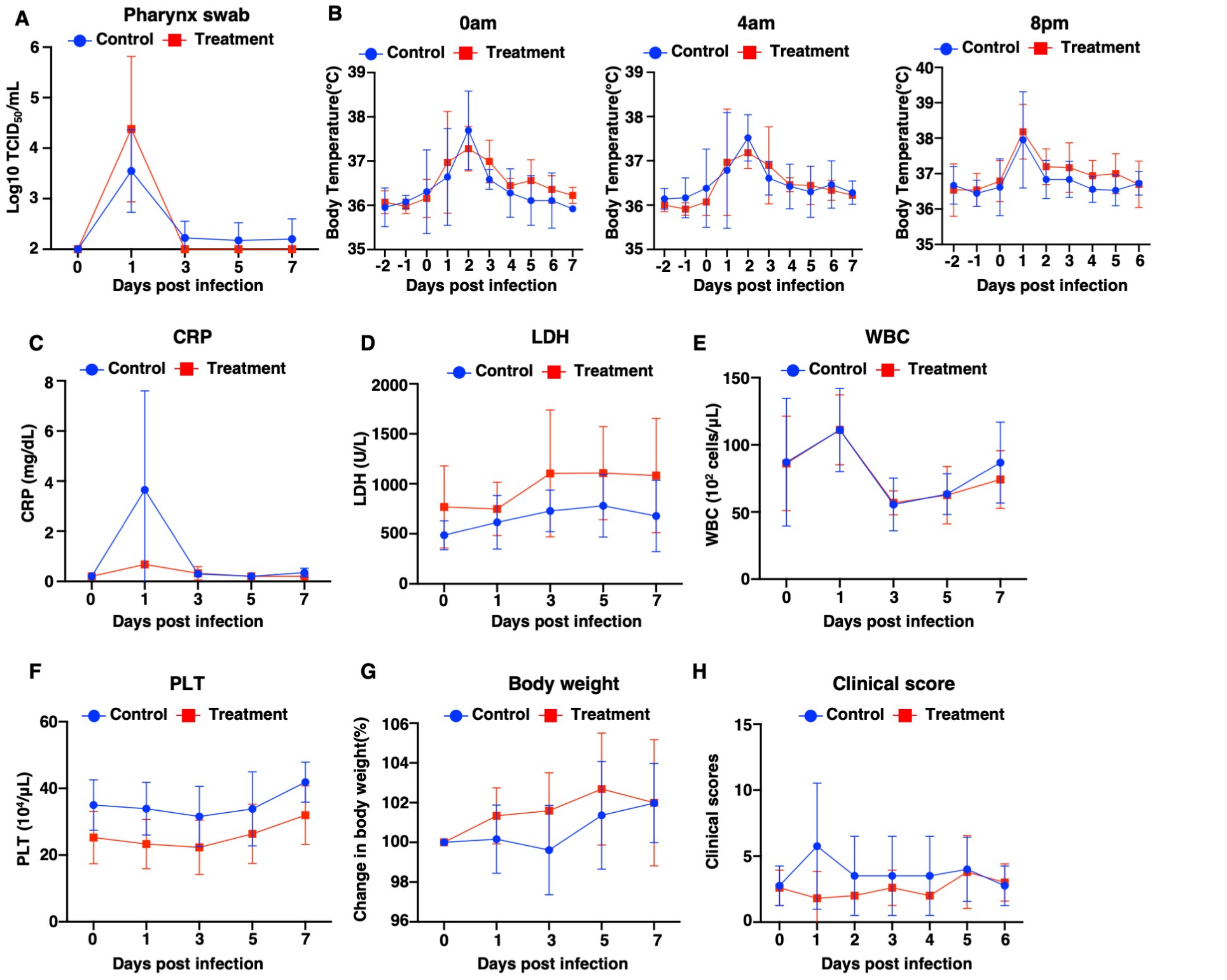


**Supplemental Figure 8. Characterization of cynomolgus monkeys infected with SARS-CoV-2.**

**(A)** Determination of viral shedding based on measurements of viral titers in pharynx swab samples. After inoculation of SARS-CoV-2, pharynx swab samples were collected at the indicated time points. **(B)** Body temperature was recorded using a data logger in infected monkeys. **(C-F)** The levels of CRP (C), LDH (D), white blood cell counts (E), and platelets (F) were measured at the indicated time points. **(G)** Changes in body weight were periodically monitored. **(H)** Infected animals were monitored for clinical signs and were individually scored daily in the following categories: appearance, secretion, respiration, discharging, appetite, and activity. Differences between the control and treatment groups were examined using two-tailed, unpaired Student *t*-test.
